# DNA Double-decker Ring Scaffolded Nanodisc for Self-assembly of Membrane Protein into Lipid Bilayer

**DOI:** 10.64898/2026.05.19.726119

**Authors:** Seaim Lwin Aye, Fatemeh Fadaei, Yuki Gomibuchi, Yuki Suzuki, Praneetha Sundar Prakash, Soumya Chandrasekhar, Takuo Yasunaga, Thorsten L. Schmidt, Yusuke Sato

## Abstract

Membrane models of scaffolded discoidal lipid bilayers called nanodiscs have proven to be a valuable tool for the study of membrane proteins in a native environment. DNA-scaffolded membrane model has emerged as an alternative tool for membrane protein studies. Taking advantage of the designability of DNA nanostructure, we created a double-decker double-stranded DNA ring (DDring) to self-assemble DNA-based nanodiscs (DNA-ND). The DDring is 17 nm wide and 4 nm high, and equipped with 28 alkyl chains on the inside that can interact with each hydrophobic leaflet of the lipid bilayer. We further demonstrate the functionality of DNA-ND membrane model with the assembly of membrane proteins. DDrings are suited to neutral or cationic charged phospholipids and detergents. This study provides more insights into the potential use of DNA- assisted nanodiscs for membrane protein characterization.

## INTRODUCTION

Membrane proteins (MPs) play a pivotal role in signal transduction and biomolecular mechanisms of living cells.^1^ Despite their importance, MPs present notorious challenges in proteomics due to the difficulty in purification and stabilization of MPs outside of the native membrane environments.^2^ Therefore, numerous lipid membrane mimetic systems have been developed to provide solutions to these challenges.^3^ Detergents micelles are most commonly used for membrane protein preparation and structural studies.^4^ Although detergents are easy to handle for preparing MPs, the detergent micelles have adverse effects on MP and its stability, such as incorrect MP folding,^5^ interference in protein-protein binding,^6^ and precipitation of soluble protein domains.^7^ Bicelles, a mixture of detergents and lipids, offer more native-like lipid bilayers for MPs.^8^ However, the presence of detergents can interfere with the stability of MP. The most native membrane mimetics are pure detergent-free lipid systems. Notably, such a membrane model called nanodisc (ND) has been employed as a promising tool for providing a native membrane-like environment for MPs.^9^ NDs are water-soluble monodisperse discoidal lipid patches comprised of a lipid bilayer held together by two copies of an amphipathic scaffold, that surround the hydrophobic region of the bilayer. NDs have been proven to be suitable for several membrane protein-related works, such as reconstitution,^10^ and structural studies,^11^ as well as a tool for coordinating with cell-free synthesis,^12^ cellular imaging,^13^ and delivery of therapeutics.^14^

The pioneer of the ND technology is the protein-based ND, a planar lipid bilayer scaffolded with amphipathic helical belt protein called membrane scaffold protein (MSP), a truncated version of human apolipoproteinA-I.^15^ The size of a ND depends on the length of the MSP protein construct used for assembly and ranges from 6.4 nm to 17 nm.^16^ The typical MSP construct MSP1D1 can form discs of ∼10 nm diameter with a molecular mass of ∼150 kDa.^17^ Protein-based NDs are well developed and suited for several applications in membrane protein studies,^18^ as well as membrane biochemistry and biophysics^15^. However, their large molecular weight and protein domains can interfere in some structural studies such as, NMR applications with an upper limit of ∼100 kDa.^19^ Although smaller NDs suitable for NMR can be achieved by deleting some helices of the MSP,^20^ the smaller NDs are prone to aggregation and can become unfit for stabilizing MPs of large sizes.

The second most well-known NDs are polymer-based NDs in which the scaffolds are amphipathic styrene maleic acid (SMA) copolymers.^21^ SMA can solubilize the membranes and form SMA lipid particles (SMALPs) to directly extract MP from the cell membrane. The major advantage of SMALP over MSP-ND is that they do not require detergents in MP preparation and can avoid the adverse effects from the detergents.^22^ However, the stability of SMALPs are susceptible to pH range: the experiments with SMAs require to be performed above pH 6.5-7, since at lower pHs, SMAs become insoluble because of the increased hydrophobicity.^23^ They also have binding affinity to divalent cations that can lead to the precipitation of the polymers.^24^ Both protein-scaffolded and polymer-scaffolded NDs offer their strengths and limitations as a tool in MP study.^25^ While MSP-NDs are easy to prepare and useful for the study of MP with the desired phospholipids and protein expression methods, they are limited in size and the MSP itself can interfere with the analysis and structural elucidation. Polymer-ND require native lipids and only MP from the living cell materials can be stabilized, limiting the range of MPs that can be studied. To address some of the issues, chemical modifications have led to the development of several SMA derivatives,^26^ and NDs with improved buffer compatibility.^27^ Despite these developments, the polydispersity of SMA can result in the size inhomogeneity of NDs due to the broad range of polymer chain lengths.^28^

Another tool used for MP reconstitution is saposin-derived lipid nanoparticles.^29^ Saposins are lipid-binding proteins, and Saposin A is the most widely used variant for NDs preparation and the size of ND depends on the Saposin A-lipid ratio and consequently are adaptable for large MPs. However, saposin proteins are more difficult to produce than MSP proteins and make it more suitable for cryo-EM than NMR applications. Another type of NDs are peptide-based NDs, where ApoA-I derived peptide is used for ND preparation.^30^ These peptides can directly extract MPs without using detergent,^31^ and the size of the ND can be adjusted by changing the peptide-lipid ratio and can achieve 30 nm-sized macrodisc.^32^ However, peptide-NDs are relatively less stable, and peptide constructs are more expensive than MSP proteins. Therefore, further developments in membrane models are still needed for the structural and functional studies of MPs.

Structural DNA nanotechnology is a tool for the de novo design of nanostructures in a programmable manner.^33^ It is widely recognized that DNA amphiphiles can interact with lipid membrane.^34^ Amphipathic DNA duplexes,^35^ and large DNA structures^36^ have been adapted to artificial cell systems to mimic biological structures, functions and mechanisms. The interaction between DNA nanostructures and lipid can be achieved via electrostatic interactions or by modifying DNA with hydrophobic moieties.^37^ Some examples include placing DNA nanostructures as channels across the lipid bilayer,^38^ encapsulating DNA nanocages inside the liposomes,^39^ and inserting nuclear pore complexes,^40^ or various types of DNA nanopores into the lipid bilayer.^41^ DNA nanostructures have also been integrated into the ND technology by using DNA-encircled lipid bilayer (DEB),^42^ DNA origami barrel,^43^ and DNA origami patterning.^44^ Among DNA-based NDs, DEBs applied a direct approach by selective alkylation of a dsDNA minicircle (dsMC) to create a belt for the hydrophobic rim of the lipid bilayer. Although this feature directly mimics MSP, dsMC is relatively thin compared to a typical thickness of the lipid bilayer, which can result in weak lipid-DNA binding. To improve the binding of DNA to the lipid bilayer, a simulation-based study using dsMC suggested the complete neutralization of dsMC inner rim.^45^ The current development of DNA-scaffolded ND technology is still in the early stages and requires further development for their application in the studies of MPs.

Here, we developed DNA-scaffolded membrane model in which a double-decker of double-stranded DNA (dsDNA) ring (DDring) scaffolds the hydrophobic leaflets of the lipid bilayer. We designed the DDring with the rationale that the reported DEB^42^ can impose weakness in the stability of the DNA structure itself and that of the lipid bilayer due to the employment of dsDNA duplex as the bilayer scaffold. Our aims in this report are (1) to improve the structural stability of DNA-scaffold, (2) to enhance the lipid bilayer and DNA-scaffold binding, (3) to ensure proper anchoring of a large variety of lipid types, and (4) to assess the potential application of DNA-based ND for MP incorporation. DDring also mimics the feature of MSP-based ND where two copies of MSPs are scaffolding the discoidal bilayer. We designed a circular single-stranded DNA minicircle (ssMC, 294 nucleotides (nt)) and hybridized with seven different complementary staples (42 nt each, 42 × 7 = 294 nt) to achieve DDring with the dimension of 16.7 nm diameter × 4 nm height. The inner rim of each ring is selectively alkylated to render the hydrophobicity at DNA-lipid interface. We performed similar lipid reconstitution method as in MSP-ND for lipid reconstitution except using detergent removal resin in place of bio-beads. We explored the behaviors of electrostatic and hydrophobic interaction between lipid and DNA upon changing the parameters of the assembly process such as detergent and lipid types. We then incorporated monomeric bacteriorhodopsin (bR),^46^ a seven transmembrane protein from *Halobacterium salinarum*, to assess the potential of DNA-ND as a membrane model for MP study. We chose bR as a model for the general class of seven-transmembrane receptors with extensive information on the structural, biophysical, and spectroscopic characterization. We performed bR incorporation as in MSP-ND, where a secondary detergent-solubilized bR is added to the primary detergent-lipid-DDring mixture prior to detergent removal. We anticipate that bR-incorporated DNA-ND can further be subjected to the functional and structural analysis to verify its use as a membrane model in MP technology.

## Results and Discussion

### Design and assembly of DNA double-decker ring

The workflow of DDring preparation involved three steps: (1) ssMC preparation, (2) alkylation of phosphorothioate (PS)-linked staples, and (3) the assembly of DDring (Figure 1A). Table S1 and Supplementary Figure S1 show the sequence and design map of ssMC positioned to the PS-linked oligos to achieve a DDring. Each PS-linked staple is 42 nt long and contains eight internal-PS modifications, 19% of PS-modifications per ssDNA strand. Upon folding, each inner rim is linked to 28 alkyl moieties, making up the total number of 56 alkyl moieties per DDring, which can hydrophobically interact with the fatty acid chains from each leaflet of the lipid bilayer. The final DDring of 294 base pairs (bp) possesses the height of 4 nm, the outer diameter of 16.7 nm, and the inner diameter of 14.7 nm.

**Figure 1.**
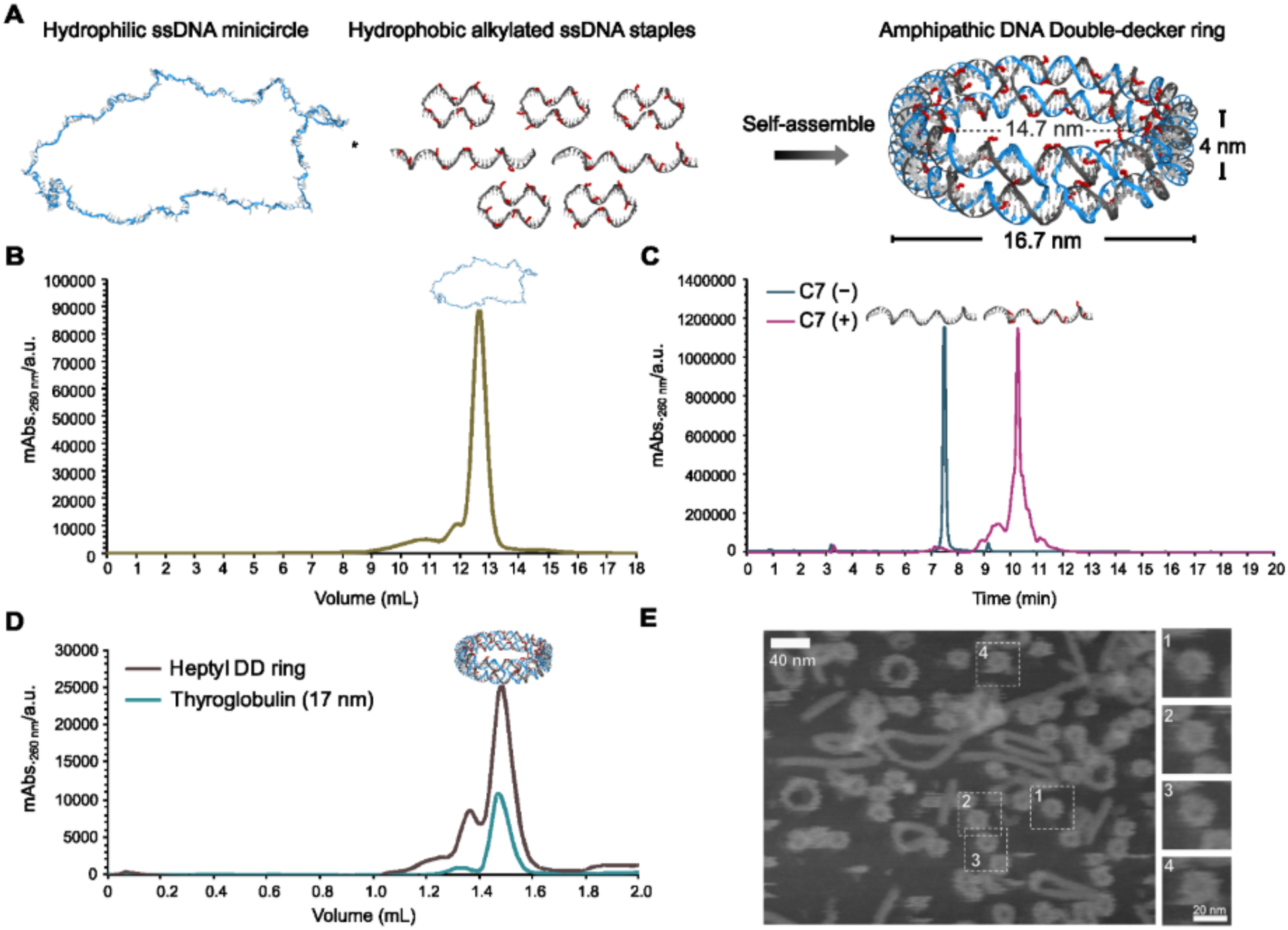
(**A**) Schematic of DNA double-decker ring (DDring) assembly. A circular ssDNA minicircle (ssMC) is annealed with seven staples modified with 8 phosphorothioates (PS) each to which alkyl moieties are attached. The alkyl moieties position inside the double-decker with the feature of membrane scaffolding hydrophobic rims. The complementary staples hybridize with the ssMC to form a dsDNA minicircle (dsMC), which in turn fold it into two decks through crossovers, ultimately forming dsDNA double-decker with the estimated dimension of ∼16.7 nm × 4 nm. (**B**) SEC profile of ssMC after exonuclease treatment. SEC was run at 0.5 mL/min elution rate with the buffer composed of 200 mM Na_2_SO_4_, 5 mM MgSO_4_ and 50 mM HEPES (pH 7.4) using Superose^TM^ 6 Increase 10/300.

We performed an enzymatic splint ligation to achieve ssMC.^47^ Two linear 147-mer ssDNAs were phosphorylated, hybridized with two complementary 30-mer ssDNAs, and ligated with T4 DNA ligase (Supplementary Figure S2). We then treated the ligated samplee with the exonucleases I and III (Exo I/III) to degrade the non-ligated linear DNA of varied lengths.

Denaturing PAGE shows that the bands representing the linear fragments before the Exo treatment in lane-1 disappeared after the Exo I/III treatment in lane-2 (Supplementary Figure S3A). The additional bands of the larger ligation products were also present in both lane-1 and lane-2. Purification by size exclusion chromatography (SEC) removed these larger products that remained after the exonuclease treatment (peak-A, Figure 1B and Supplementary Figure S3B). The peak-B from SEC represented a sharp band of the target ssMC in lane-3. Although SEC increased the purity of ssMC, the yield was very low (∼20%), leading us to perform only exonuclease digestion for most of the ssMC preparation.

To create amphipathic DDring, the hydrophilic ssMC hybridizes with the hydrophobic staples. To render hydrophobicity to DNA, we modified the seven PS-linked staples with alkyl moieties^48^ by designing the sequences with 28 curved A-tracts to strategically position the alkyl chains around the inner rims of DDring.^49^ After alkylation, we separated the samples based on their polarity by reverse-phase high-performance liquid chromatography (RP-HPLC) using C18 column. The RP-HPLC chromatogram of heptyl (C7) staples in Figure 1C shows the change in retention time from the starting material C7(−) to the C7(+) staples. Although we also alkylated PS-linked staples with Iododecane (C10), the yield of the C10(+) staples was extremely low due to the lower solubility in water and higher adsorption to the column (Supplementary Figure S4). The denaturing-PAGE after RP-HPLC purification also confirmed the hydrophobicity of the C7(+) staples, where C7(−) DNA migrated faster than the hydrophobic C7(+) DNA (Supplementary Figure S5A, B).

To assemble DDring, we performed the thermal annealing of ssMC after exonuclease treatment with the excess number of staples and analyzed the size of the DDring with SEC using a 10/300 GL column (Supplementary Figure S6A). Native PAGE of the SEC eluted fractions shows that, the peak-2 was the target DDring and the peak-1 and peak-4 were the larger byproducts and unbound staples, respectively (Supplementary Figure S6A, B). We used 30 kDa molecular weight cut-off (MWCO) filter to remove the excess staples (Supplementary Figure S6C). Higher molar ratios of the staples to ssMC generally gave the more well-formed DDring (Supplementary Figure S7A). The comparison of ssMC and DDring shows the slower migration through native PAGE gel with dsDNA DDring, confirming the successful DDring formation (Supplementary Figure 7B). SEC elution profile also confirms that the size of the purified DDring_C7(+)_ overlapped with the retention volume of the standard protein, Thyroglobulin (Stokes diameter of 17 nm) (Figure 1D). Since DDring was prepared from ssMC after exonuclease treatment, the larger byproducts were also present in the earlier elution, as observed in the SEC chromatogram of ssMC. Atomic force microscopy (AFM) imaging further confirmed these results with the presence of larger byproducts in addition to the targeted DDring (Figure 1E).

The remaining non-targeted circularized DNAs that are not digested with exonucleases, including dimers were eluted from 10 to 13 mL. The ssDNA scaffold was eluted from 13 to 14 mL. The eluted fractions were analyzed in denaturing PAGE that is shown in Supplementary Figure S3. (**C**) RP-HPLC profiles of DNA before and after alkylation. DNAs with 8 PS groups were treated with iodoheptane at 65 °C for 2 h. DNA without heptyl side chains (C7−) was eluted at 8 min and DNA with heptyl side chains (C7+) was eluted around 11 min. (**D**) Size exclusion chromatography (SEC) profile of heptyl DDring in comparison to the protein marker, Thyroglobulin, with Stokes diameter of 17 nm. SEC was run with Superose^TM^ 6 Increase 3.2/300 at a flow rate of 0.04 mL/min. The elution buffer is composed of 150 mM NaCl, 5 mM MgCl_2_ and 20 mM HEPES (pH 7.4). (**E**) AFM images of DDring, scale bar = 40 nm.

### Lipid bilayer formation within DDring

The schematic in Figure 2A illustrates the process of the lipid bilayer reconstitution within DDring. The general procedure of DNA-ND preparation is as follows: the detergent-lipid solution was mixed with the DDring solution in a 1:450 molar ratio and incubated for 1 h at the temperature appropriate for the lipid. Then, the gradual detergent removal with the resin was carried out over a 4–6 h period guiding the self-assembly of the lipid bilayer within DDring. The phospholipid mixture was mainly composed of ∼70–100% zwitterionic lipids, namely DMPC or POPC, 10–30% cationic lipids, either DMTAP or DOTAP, and 1–2% of fluorophore-labeled lipid, Rhodamine-PE. The buffer solution contained Mg^2+^ and Na^+^ ions to decrease the electrostatic repulsion of the unmasked DNA with lipid.^50^ To assess the density of the lipid and DNA after the detergent removal, we used the overlaid Iodixanol density gradient ultracentrifugation (DGUC).

**Figure 2.**
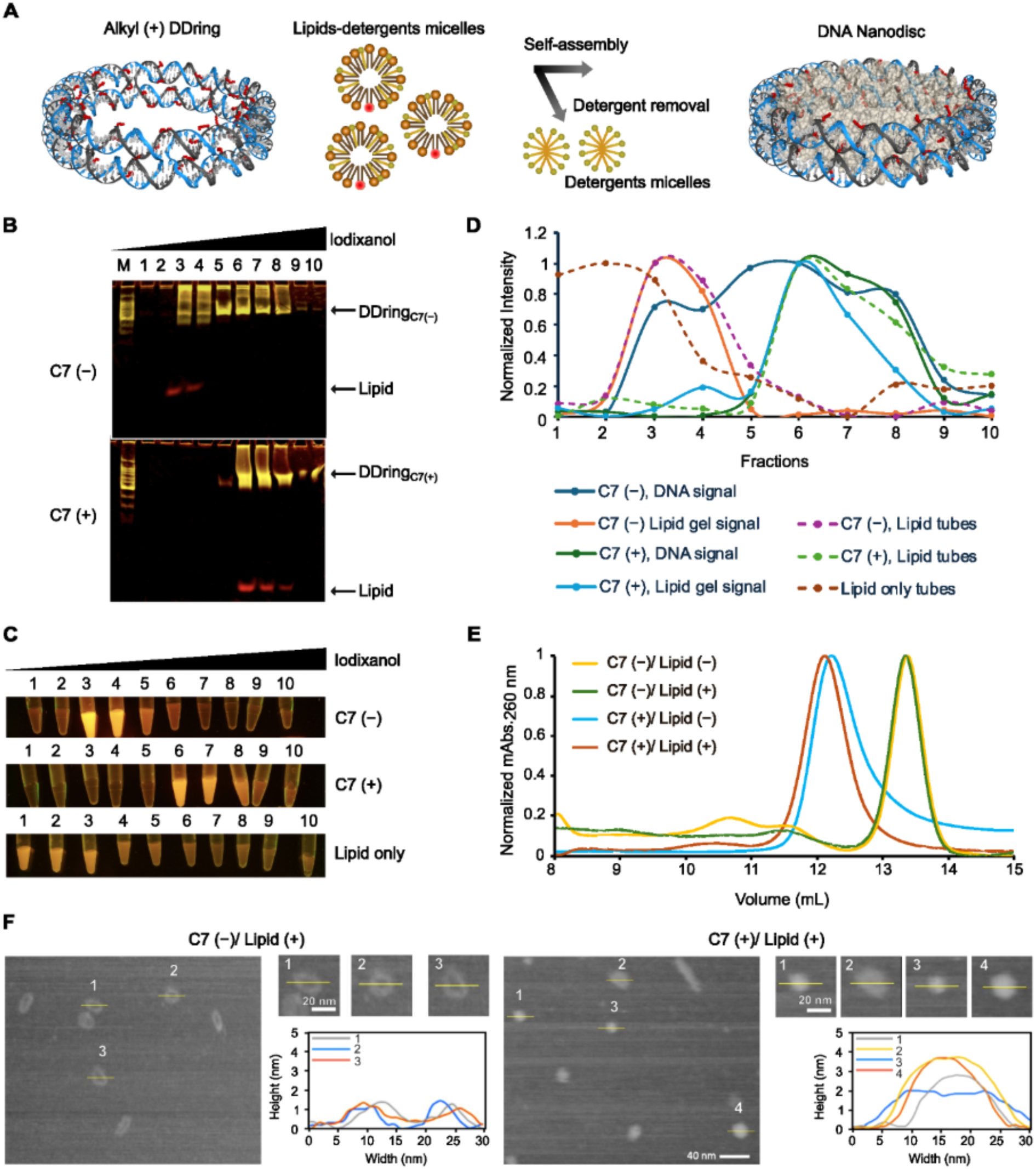
(A) Schematic of DNA-nanodisc reconstitution. For a general procedure, DDrings are mixed with lipids solubilized in detergents, incubated for a brief period, and the detergent is removed to complete the lipid bilayer assembly within DDring. (**B**) Density gradient ultracentrifugation of Rhodamine-PE-containing lipids mixed with DDrings with and without 58 heptyl moieties (C7 (−/+)). Fractions (1–10) were taken by pipetting from top to bottom, with the ascending Iodixanol concentration and thereby the increasing density and analyzed with 5% SDS-PAGE supplemented with 100 mM NaCl. M = 100 bp ladder. Lipid = POPC:DOTAP:Rhodamin-PE (69:30:1), detergent = 16 mM CHAPS, DNA: Lipid = 1:450. (**C**) Normalized intensity of lipids and DDring from B. Lipid only image is shown in Supplementary Figure 10. (**D**) Normalized elution profile of a size exclusion chromatography (SEC) before and after lipid reconstitution within (C7 (−/+)) DDring. DNA-lipid reconstitution concertation is the same as in (B). (**E**) AFM image of DDring with and without heptyl moieties (C7 (−/+)) after lipid reconstitution with their height profiles.

The choice of the detergent plays a significant role in MSP-based ND reconstitution, despite that the detergent is removed in the final formed ND.^51^ Similarly, the tested detergents with varied characteristics, anionic sodium cholate, zwitterionic CHAPS and cationic DTAB, behave differently in DNA-based ND. In case of sodium cholate-solubilized DMPC/DMTAP, the profile of normalized intensity from PAGE analysis after DGUC shows that both lipid and DNA were present in the denser fractions of DDring_C7(+)_ (Supplementary Figure S8A). However, with sodium cholate-DMPC/ DMTAP, the electrostatic interactions between hydrophilic DDring_C7(−)_ and the lipids was also prominent, rendering the inconclusive comparison between DDring_C7(−)_ and DDring_C7(+)_. Next, to exclude the negative charge from the detergent, we replaced sodium cholate with CHAPS, while the other parameters remained the same.

Normalized DGUC data indicated a similar lipid-DNA interaction as in sodium cholate, with a more distinct colocalization pattern between DDring_C7(+)_ and the DMPC/DMTAP, indicating that the choice of the detergent can alter the assembly result (Supplementary Figure S8B).

The thickness of DMPC/DMTAP bilayer is relatively short compared to the height of DDring,^52^ and this can contribute the stronger electrostatic interaction with the hydrophilic DDring and the lipid head groups. The change of the temperature after detergent removal at 25 ℃ to DGUC at 4 ℃ can alter the fluid state (*L_⍺_*) of DMPC bilayer phase to the sub-gel state (*L_c_*) and the lipid-DNA interaction can become more prominent. To test this hypothesis, we changed the lipid composition to the CHAPS-solubilized POPC/DOTAP mixture and showed the prominent difference in the DNA-lipid colocalization between DDring_C7(+)_ and DDring_C7(−)_ (Figure 2B, C). DDring_C7(−)_ overlapped with the lipids only to a small extent in the third and fourth fractions, while the lipid and DDring_C7(+)_ colocalized at the denser fractions. In the lipid only experiment without DNA, lipid after detergent removal dispersed in almost all fractions starting from the first one (Figure 2C, Supplementary Figure S9A, B). The POPC/DOTAP bilayer was also tested with DTAB, which showed a similar DNA-lipid colocalization pattern to CHAPS (Supplementary Figure S9A–D). These results suggested that different types of lipid bilayer and detergent can affect the reconstitution process. Zwitterionic detergent CHAPS performed very well with zwitterionic lipids, DMPC and POPC. Cationic detergent DTAB performed better than anionic detergent Na-cholate. DMPC/DMTAP combination showed more electrostatic interaction with hydrophilic DDring compared to POPC/DOTAP.

Next, we compared the size of the DDrings with and without POPC/DOTAP reconstitution by SEC (Figure 2D). The SEC profile indicated a slightly larger size with DDring_C7(+)_ compared to DDring_C7(−)_, as expected. Due to the rigid structure of DDring, the lipids should fill in the inner space, without significantly transforming its shape and size. Therefore, the SEC profile shows a slight change in the retention volume of DDring_C7(+)_ with lipids. To confirm the disc formation within DDring_C7(+),_ DGUC purified DNA-fraction was subjected to AFM imaging (Figure 2E and Supplementary Figure S10A–C). DDring_C7(−)_ with lipids appeared empty, while DDring_C7(+)_ revealed a filled disc feature. AFM analysis of DDring_C7(−)_ revealed the average height of ∼2 nm that is lower than the actual height of 4 nm due to the sample deformation caused by the AFM tip-sample interaction.^53^ The heights of the lipid-filled DDring_C7(+)_, DNA-ND are about 4 nm, suggesting the presence of lipids within the rings.

### Self-assembly of Integral Membrane Protein, Bacteriorhodopsin in DNA-ND

We chose bacteriorhodopsin (bR) as a target protein for incorporation into DNA-ND. bR is a membrane protein with seven transmembrane helices from *Halobacterium salinarum*. A single bR is ∼4.3 (± 5 nm) in diameter,^54^ with a surface negative charge,^55^ and molecular weight of ∼27 kDa.^56^ bR incorporates efficiently into protein-based NDs as a monomer (70–90%),^57^ as well as trimer.^58^ bR can be solubilized in monomeric form in both Triton X-100 and n-octyl-β-D-glucoside (OTG).^59^ We selected the mild non-ionic detergent OTG for bR solubilization due to its higher critical micelle concentration than Triton X-100.

The schematic in Figure 3A demonstrates the incorporation of bR within DNA-ND. As stated above, we used stoichiometric ratio of 1:450 (DDring: lipids) for empty DNA-ND reconstitution. Due to the displacement of lipids by bR, we reduced the lipid number to 400 since about 60 of POPC lipids and 70 of DMPC can give the bilayer of 4.4 nm.^60^ The space of the reduced lipids should be able to accommodate ∼4.3 nm bR monomer. bR incorporation process is the same as the empty DNA-ND reconstitution. DDring solution was first mixed with the lipid solubilized in the primary detergent and incubated for 30 min, followed by the addition of the secondary detergent-solubilized bR solution. After incubating the DNA-lipid-protein mixture for another 30 min, the two detergents were gradually removed over a 4–6 h period. We purified the detergent-free samples with SEC by monitoring the absorbance at 260 nm that detects the DNA signal. Due to the extremely low amount of bR (0.1 μM) in the sample injected into SEC, absorbance at 550 nm was not detectable. We eluted four individual peaks for DDring_C7(+)_, 1, 2, 3 and 4, and two individual peaks for DDring_C7(−)_, 1 and 2, from the respective SEC chromatograms (C7_(+)_ and C7_(−)_, Figure 3B). By running two PAGE analysis, one for DNA and another for protein in parallel, we compared DNA and protein signals in the elution profiles. The detection of both DNA and protein bands in lane-4 (C7_(+)_) revealed that DDring_C7(+)_-based ND contained bR, whereas bR was not detectable in lane-2 (C7_(−)_) (Figure 3C). We repeated bR-ND assembly with DDring_C7(+)_ using both POPC/DOTAP and DMPC in parallel and purified with DGUC. DNA fractions in SDS-PAGE were pooled and analyzed again for both protein and DNA detection. Both POPC/DOTAP and DMPC batches showed that the lipids, proteins and DNA were present in the pooled collection, confirming the SEC result (Figure 3D). We also studied bR-incorporated DNA-ND by negatively stained transmission electron microscopy (TEM) (Figure 3E). The comparison between with and without bR batches showed the presence of bR with a different appearance to the empty disc, indicating the bR incorporation within DNA-ND.

**Figure 3.**
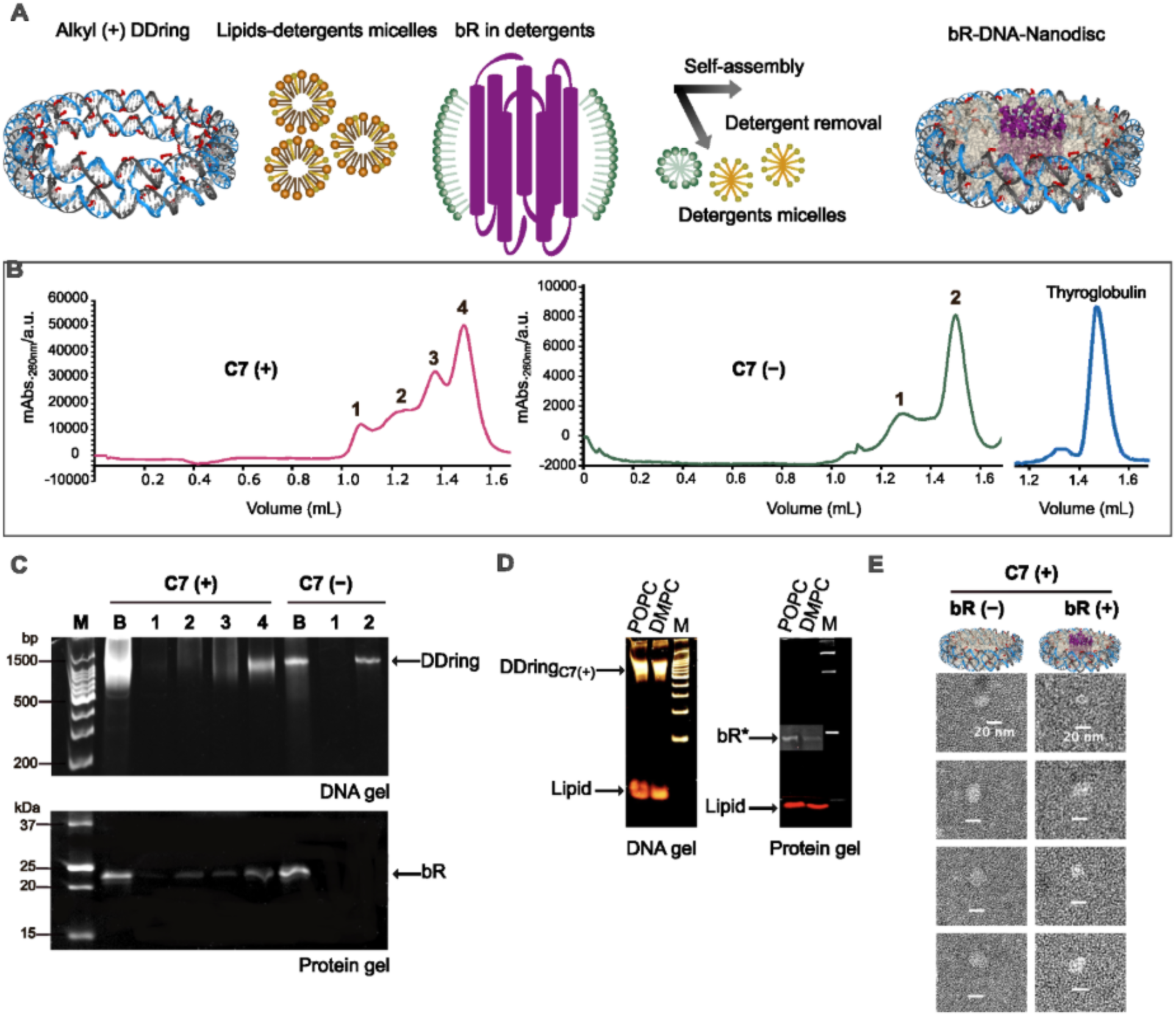
(**A**) Schematic of bacteriorhodopsin (bR) incorporation into DDring-scaffolded nanodisc, not scale to the actual size. DDrings are first mixed with lipids in detergents followed by the addition of membrane protein solubilized in a secondary detergent. After incubating for a brief period, the detergents are removed to assemble the lipid bilayer and protein within DDring. (b) SEC profiles of assembly mixture C7 (−/+) DDring with DMPC and bR. Protein standard, thyroglobulin (Stokes diameter = 17 nm). DMPC lipids were first dissolved in the buffer with CHAPS and bR with Octyl-β-D-thioglucoside. DDring:Lipid = 1:400, DDring: bR = 2:1, Buffer = 10 mM HEPES (pH 7.4), 5 mM MgCl_2_. Superose 3.2/300 was used for SEC analysis and run with a flow rate of 0.04 mL/ min. The eluted fractions from SEC chromatogram were marked with number s 1–4 for C7 (+) and 1 and 2 for C7 (−). (**C**) SDS-PAGE analysis of the eluted fractions for DNA was done with 6% gel containing 100 mM NaCl and protein analysis with 15% gel. B = the assembly mixture after detergent removal and before SEC elution. Marker (M) for DNA = 100 bp ladder and marker for protein = Precision plus protein standards. (**D**) Density gradient ultracentrifugation analysis of DNA-ND assembly after reconstitution C7(+) DDring with lipids and bR. Fractions containing DNAs were pooled and concentrated with 10 kDa molecular cut off filter and further analyzed in SDS-PAGE for protein and DNA. POPC = POPC:DOTAP:Rhod-PE (83:15:2) and DMPC = DMPC:Rhod-PE (98:2). (**E**) Representative negative stain TEM images of DMPC nanodisc with and without bR.

### Atomistic Molecular Dynamics Simulations

To examine the stability of heptyl-functionalized DNA nanodiscs, we carried out all-atom molecular dynamics simulations over 200 ns using two different lipid compositions, DMPC/DMTAP and POPC/DOTAP. In both cases, the DNA scaffold remained intact and stably attached to the lipid bilayer throughout the simulation, with no signs of detachment or structural breakdown. This stable association is driven by the heptyl modifications, which insert into the hydrophobic core of the lipid bilayer and effectively anchor the DNA nanodisc to the membrane. As shown in Figure 4, this configuration was established from the start of the simulation and was consistently maintained at 100 and 200 ns, regardless of the lipid composition used. Movies 1–4 show side and top views of DMPC/DMTAP and POPC/DOTAP, highlighting the stable association of the lipids with the nanodisc. Taken together, these results show that heptyl functionalization is a reliable and effective way to attach DNA nanodiscs to lipid bilayers, and that the resulting structures are stable enough to support their use in membrane-related applications. Furthermore, the close-up from the last frame shows that the alkyl chain anchors to the lipid, causing the DNA nanodisc and lipids to stick together, confirming that the DNA nanodisc remains stable.

**Figure 4.**
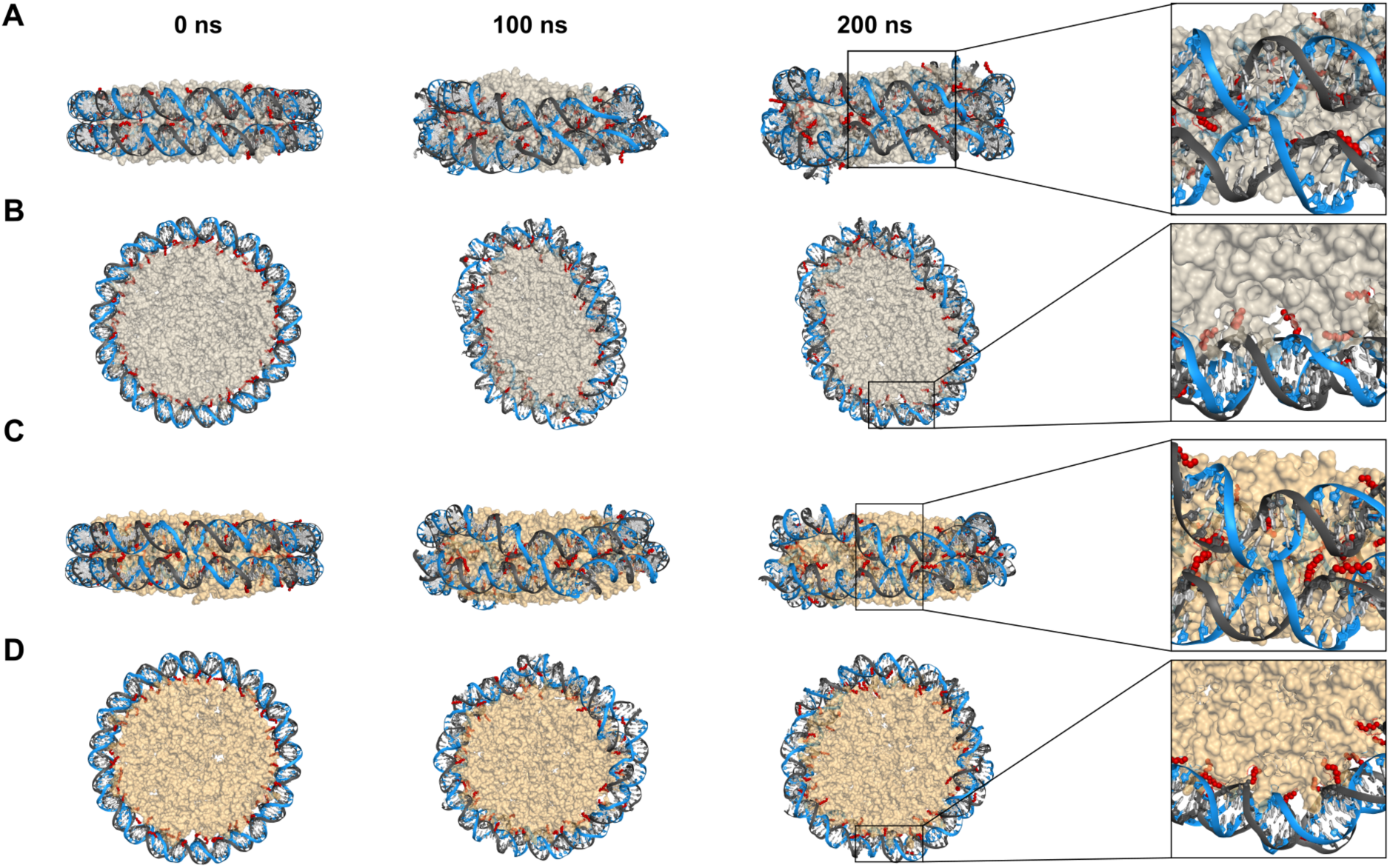
All-atom molecular dynamics simulations of heptyl-functionalized DNA nanodiscs over 200 ns. **(A)** Side and **(B)** top views of DNA nanodiscs with DMPC/DMTAP lipids shown as a light gray surface. **(C)** Side and **(D)** top views of DNA nanodiscs with POPC/DOTAP lipids shown as a light orange surface. In both systems, the heptyl-modified DNA strands (dark gray) are inserted into the lipid bilayer through heptyl groups (red spheres), anchoring the DNA scaffold to the nanodisc. Insets show magnified views of the lipid–DNA interface at 200 ns, highlighting the stable intercalation of heptyl chains into the hydrophobic core of the lipid bilayer. Snapshots are shown at 0, 100, and 200 ns.

**Figure 5.**
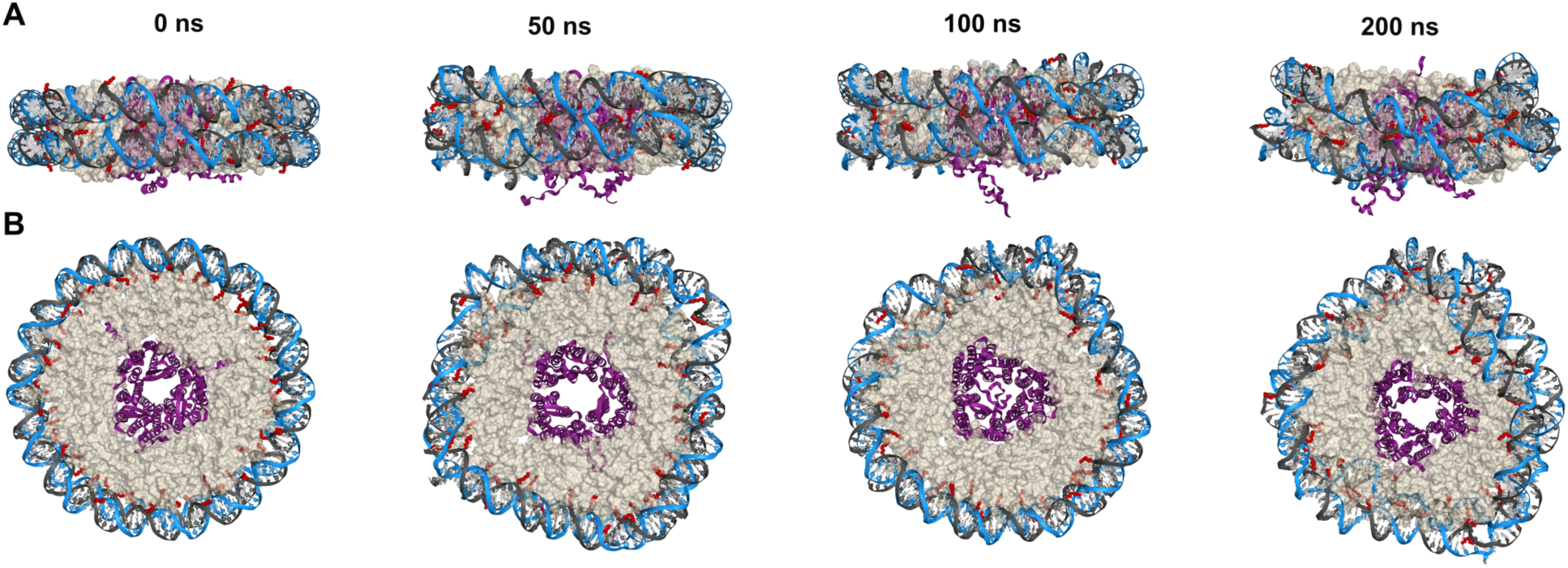
All-atom molecular dynamics simulation of bR trimers embedded in DNA nanodiscs over 200 ns. (A) Side and (B) top views of the DNA nanodiscs. The DMPC/DMTAP lipid environment is shown as a light gray surface, with heptyl groups represented as red spheres. The DNA scaffold is shown in light blue, heptyl-modified DNA strands in dark gray, and the membrane protein (bR) in purple. Snapshots are provided at 0, 50, 100, and 200 ns to illustrate the stability of the system.

Since the DNA nanodiscs remained stable in both lipid combinations (DMPC/DMTAP and POPC/DOTAP), we proceeded to incorporate membrane proteins into the discs. We successfully demonstrated that bR trimers can be embedded into the DNA nanodisc. Over the course of a 200 ns MD simulation, the protein demonstrated stable integration, remaining bound to the DNA nanodisc structure throughout the trajectory. Movies 5 and 6 show the MD trajectories from the side and top perspectives, respectively, highlighting the stable association of the bR trimer with the nanodisc.

## Conclusion

In this report, we undertook the engineering of an amphipathic DDring that encircles the discoidal lipid bilayer, resembling the feature of two scaffolding proteins in protein-based NDs. Although the previous DEB reported to be able to reconstitute DMPC bilayer within dsMC,^42^ it employed one dsMC helix with 2 nm thickness, which may not be enough to accommodate bilayer with longer hydrophobic chains such as POPC. In comparison, the height of DDring was doubled to 4 nm and contain two times more hydrophobic moieties, rendering the conceptually more stable lipid-DNA binding. DDring is made of two interconnected DNA minicircles, in which alkylated staples folded one long circular DNA scaffold (ssMC) into the shape of two dsDNA rings stacked on top of each other. We found that the circular DNA byproducts of non-targeted size from the ssMC preparation can decrease the homogeneity of DDring size. In future, this issue can be addressed by using a single linear ssDNA as starting material or by adapting different ssMC preparation.

Since the number of PS modifications per oligo is limited, we selectively alkylated DDring with ∼20% per staple, incorporating 56 alkyl moieties around the ∼15 nm inner rims of double-decker. The previous dsMC used a single staple attached with four butyl or decyl moieties to hybridize with the repetitive sequences along ssMC.^42^ However, the choice of decyl moieties is not feasible with DDring. Instead of repetitive sequences, we used seven different staples with 8 PS groups and modifying them with decyl moieties increased the synthesis complexity and aggregation. Therefore, we used the heptyl moieties for the alkylation of seven staples to achieve the amphipathic DDring.

DNA-ND assembly reported here is similar to that of protein-based ND and requires the solubilization of the phospholipids with detergent. Instead of Bio-beads or dialysis, we used detergent removal resin that is available for the sample of μL scale. Although it is unknown that the quick removal of detergent can hinder the lipid assembly, we modified the manufacturer’s protocol and extended the detergent removal period to several hours. We also explored the parameters in the self-assembly of lipid bilayers and found that the choice of detergent and phospholipid can affect the lipid assembly. Zwitterionic detergent with the higher critical micelle concentration performed the best compared to charged detergents, while cationic detergent performed better than anionic detergent. We also found that, when the cationic lipids were present, their headgroups could interact via electrostatic interaction with the DDring without hydrophobic modification. The electrostatic interaction was more prominent with DMPC than POPC.

It is crucial for DNA-ND to function as a native-like discoidal membrane model for MP. DNA-ND can become a valuable tool for the structural elucidation of MP due to the lack of extra protein. We incorporated bR in the assembly mixture to determine whether the detergent-solubilized protein can be incorporated into DNA-ND. Both POPC and DMPC bilayers can incorporate bR into DDring, without the cationic lipid in DMPC case. Although we investigated bR incorporation with DGUC, SEC, and TEM, the bR orientation as well as the structural and functional integrity of bR in DNA-ND need to be explored further. Compared to protein-based ND, the development of DNA-based ND is still at an early stage. The challenge for DNA-scaffolded ND lies in the successful hydrophobic modification. Although the alkylation is economical, the extended period of heating, and repeated drying can lead to the DNA degradation, which can disrupt the correct stoichiometry of DNA, lipid, and protein. Therefore, alternative approaches for hydrophobic modification should be explored for DNA scaffolding.

The programmable DNA nanotechnology will allow the design of DDrings with diverse sizes and shapes that can meet the requirement imposed by the target MP. DNA-ND can be linked with larger DNA origami structures, which can potentially be useful in studying protein-protein interaction by arranging ND with different MP side by side. We demonstrated here that bR without any modification can be incorporated into DNA-ND. In the future development, MP itself can be modified with ssDNA tags or linker complementary to the linker from DNA-scaffold for the desired purposes, such as to control the orientation of MP or to study protein-protein interaction. We anticipate that DDring design reported here will be a promising tool to construct the lipid membrane model for the study of MP and other biological applications.

## METHODS

### Alkylation of phosphorothioate oligonucleotides

A typical method for the alkylation of PS-linked oligos is as follows: commercially modified PS-linked oligos (Eurofins, HPLC grade) were dissolved in UltraPure^TM^ distilled water (Invitrogen) to obtain a 100 μM stock solution. The seven PS-oligos, com-1, -2, -3, -4, -Up and -Down (Supplementary Table S1), (5 μL, 100 μM) were dried in 1.5 mL microcentrifuge tubes using Eppendorf Concentrator plus^TM^ and resuspended in Tris-HCl (4 μL, 30 mM, pH 8.0, NIPPON GENE). Then, N, N-dimethyl formamide (DMF) (36 μL, FUJIFIM Wako) and Iodoheptane (10 μL, Sigma-Aldrich) were added to the tube. The tube was heated to 65 °C for 3 h while shaking at 1, 000 r.p.m using a dry block bath shaker (Front Lab, AS ONE). It should be noted that multiple tubes were prepared in parallel, and total volume may vary while keeping the final composition of 30 mM Tris to 10% and DMF to 90%. Due to the evaporation, Iodoheptane was supplemented every hour throughout the incubation. After incubation, the reaction mixture was evaporated in the concentrator at 60 °C, dissolved in Ethylenediaminetetraacetic Acid (EDTA, 50–100 μL, 0.1 M, pH 8.0, NIPPON GENE) and then heated to 90 °C for 5 min. The samples were purified with NAP-25 columns (Cytiva) according to the manufacturer’s method. DNA-containing fractions were determined with a Nanodrop spectrophotometer (DeNovix, DS11 Fx) measuring the absorbance at 260 nm. The DNA fractions were combined and then dried with the concentrator for further purification by reverse-phase high-performance lipid chromatography (RP-HPLC).

### RP-HPLC purification of alkylated DNA

The alkyl staples after NAP purification were resuspended in 100 μL Triethylamine Acetate (TEAA) (20 mM, pH 8.0, Thermo Fisher). The RP-HPLC was carried out by using JASCO (PU-4180, AS-4050, UV 4075, BS-4000 and LC-NetII ADC) apparatuses and C-18 column (Agilent) with the flow rate of 1 mL/ min using the following gradient: 0–1 min: 3% Acetonitrile (ACN), 1.1–20 min: 90% ACN, 20.1–21 min: 100% ACN, 20.1–26 min: 100% ACN, 26.1–31 min: 100% TEAA, 31–40 min: 100% TEAA. The eluted samples were dried in the concentrator, resuspended in distilled water, and stored at −20 °C.

### SSMC preparation

Two long ssDNA sequences, 147-1 and 2, and two short sequences, Splint-1 and 2, were designed through OligoAnalyzer tool (idtdna.com) avoiding the potential secondary structure formation. Two long DNAs, 147-1 and 2 (HPLC grade), and two splint oligos (Analytical grade) were purchased from Eurofins. Phosphorylation of long oligos was performed as follows: 147-1, 2 (1 μL each, 100 μM) were mixed with 2 μL of 10× T4 DNA ligase buffer (New England Biolabs (NEB)), and 10 units (U) of T4 polynucleotide kinase (NEB) in a 20 μL reaction volume and incubated at 37 °C for 30 min. The phosphorylated solution was mixed with Splint 1 and 2 (1.5 μL each, 100 μM), 10× T4 DNA ligase buffer (2 μL), adjusting to the final volume of 40 μL with distilled water. The oligos were hybridized by using a MiniAmp^TM^ thermal cycler (Thermo Fisher Scientific) with a program of 80 °C for 2 min and gradual cooling of temperature from 80 °C to 20 °C with 5 °C/ min. The annealed mixture was then supplemented with 10× T4 DNA ligase buffer (1 μL) and T4 DNA ligase (1 μL, 400U/μL) adjusting the final volume to 50 μL. The ligation mixture was incubated for 1 h at 25 °C. The removal of enzyme and buffer was performed by using oligo clean and concentrator (Zymo Research) according to the manufacturer’s protocol. The non-ligated linear DNAs were degraded with Exonuclease I and III (NEB) as per the manufacturer’s protocol and consecutively purified by Zymo kit and kept at −20 °C until further use.

### DNA double-decker ring preparation

A typical reaction of DNA double-decker ring (DDring) preparation is as follows. ssMC and alkyl (−/+) staples were mixed in 1:5 molar ratio with 1× TE buffer (1 mM EDTA, 10 mM Tris, pH 8.0) and 8 mM MgCl_2_. The DDring folding was performed in the thermal cycler with a program of 85 °C for 2 min and cooling down to 20 °C at a rate of 1 °C/ 12 sec. The annealed product was concentrated with 30 kDa MWCO, Amicon Ultra (Millipore) filter according to the manufacturer’s protocol. The buffer was exchanged for the reconstitution buffer (200 mM Na_2_SO_4_, 5 mM MgSO_4_, 50 mM HEPES pH 7.4) using the column.

### Denaturing and native polyacrylamide gel electrophoresis

The ssMC and alkylated staples were analyzed by 5–6% and 16% denaturing polyacrylamide gel electrophoresis (PAGE), respectively. The urea-supplemented gel was prepared as follows: for 10 mL volume, 4.8 g urea (FUJIFIM Wako) was dissolved in 1.25 mL or 4 mL of 40% acrylamide/bis solution (19:1) (Nacalai Tesque), 1× Tris-Borate-EDTA buffer (TBE) (Takara Bio), 0.1% ammonium persulfate (APS) (FUJIFIM Wako) and 0.1% N,N,N’,N’-tetramethylethlyene-diamine (TEMED) (FUJIFIM Wako). The electrophoresis was performed with ATTO AE-6530 miniPAGE chamber with a gel size: 90 × 80 × 1 mm and 500 mL of 1× TBE buffer. The gel was pre-run at 45 °C, 200 V for 30 min and washed the well by pipetting. Then, 20 ng of the samples and DNA ladders (Takara Bio) were mixed with 2× loading buffer (95% formamide, 18 mM EDTA, and 0.1% bromophenol blue) and loaded to the well. The loaded gels were then run for 1 h. DDring was analyzed by 5–6% native PAGE analysis. The gel was prepared as follows: the components are added in this order, 5–6% of 40% Acrylamide/ Bis solution (29:1) (Nacalai Tesque), 1× TBE, 100 mM NaCl, 0.1% TEMED and 1% APS. The running buffer 1× TBE contained 100 mM NaCl. The marker and the sample DNA were loaded at 20 ng mixed with 6× loading buffer (5 mM Tris HCl, 50% glycerol, 1 mM EDTA, 8 mM MgCl_2_, 0.25% bromophenol blue). The electrophoresis was carried out at 4 °C by running at 100 V for 1 h. The gel was post-stained with 1× SYBR^TM^ Gold Nucleic Acid Gel Stain (Invitrogen) in 100 mL 1× TBE for 15 min and the DNA was visualized with FastGene FAS-DIGI Compat–UV-free gel imager with a built-in high-quality camera (Canon 250D, Nippon Genetics) or iBright^TM^ FL1500 imaging system (Thermo Fisher Scientific).

### Formation of DNA-Nanodisc

The phospholipid, 1-palmitoyl-2-oleoyl-glycero-3-phosphocholine (POPC) was purchased from NOF America Corporation. The other phospholipids,1,2-dioleoyl-3-trimethyl ammonium-propane (chloride salt), (18:1 TAP, DOTAP), 1,2-dimyristoyl-sn-glycero-3-phosphocholine (14:0 PC, DMPC), 1,2-ditetradecanoyl-3-trimethyl ammonium-propane (chloride salt), (14:0 TAP, DMTAP), and 1,2-dipalmitoyl-sn-glyero-3-phosphoethanolamine-N-(lissamine rhodamine B sulfonyl) (16:0 Liss Rhod PE) with (excitation/emission - 560 nm/ 583 nm) were purchased from Avanti Research. Phospholipids of the required amount were dried under the stream of nitrogen gas and then kept in a desiccator to remove the moisture. The dried lipids (2 mM) were solubilized in 200 μL buffer (200 mM Na_2_SO_4_ and 5 mM MgSO_4_, 50 mM HEPES pH 7.4) with the required detergent. The lipid solution was sonicated in a water bath sonicator (Branson M1800/CPX-952-116R) for 2 h. For lipid and DNA reconstitution, DDrings were mixed with lipid-detergent solution in a 1:450 molar ratio (for example: 300 nM DNA:135 μM lipids in 50–100 μL volume). The final concentration of the selective detergent was kept over CMC with 18 mM for sodium cholate (FUJIFIM Wako), 16 mM for 3-((3-cholamidopropyl) dimethylammonio)-1-propanesulfonate (CHAPS, Dojindo) and 30 mM for Dodecyltrimethylammonium Bromide (DTAB, TCI). The lipid-detergent-DDring mixture was incubated at the temperature appropriate for the lipids, 4 °C for POPC and 25 °C for DMPC) for 1 h. The detergent removal using Pierce^TM^ detergent removal resin and (500 μL) spin column was performed according to the manufacturer’s protocol with a few modifications as follows: The same volume of the resin slurry washed with the buffer (200 mM Na_2_SO_4_ and 5 mM MgSO_4_, 50 mM HEPES pH 7.4) was added to the DNA-lipid-detergent mixture and incubated on an end-to-end rotatory shaker for 4 h or overnight. The mixture was transferred to the spin column with the washed and dried resin and collected the detergent free flow-through of lipid-DNA mixture and stored at 4 °C .

### Protein incorporation into DNA-Nanodisc

The purple membrane protein, bacteriorhodopsin, was purchased from Sigma and solubilized by dissolving in the buffer (200 mM Na_2_SO_4_ and 5 mM MgSO_4_, 50 mM HEPES pH 7.4) and 46 mM of the detergent n-octyl-β-D-thioglucoside (OTG, Dojindo). The bR solution was sonicated for a few min and kept in the dark for 1–2 days. The protein solution was then centrifuged at 45 000 g for 10 s; the supernatant was taken into aliquots and kept at −80 °C. For a typical experiment, DDring were mixed with lipids-CHAPS mixture in (1:400) molar ratio and incubated at required temperature for the lipids. After 30 min, the bR-OTG mixture was added in the ratio of 2 DDring per protein. After 30 min, the detergent removal process was performed as in empty nanodisc preparation and kept at 4 °C for further analysis.

### Density gradient ultracentrifugation and SDS-PAGE analysis

The gradient was performed with Iodixanol solution (OpiPrep^TM^) prepared as 45%, 26%, 22%, 18%, 14%, 10%, 6%, and 2% in the buffer (200 mM Na_2_SO_4_ and 5 mM MgSO_4_, 50 mM HEPES pH 7.4). The overlay method was used for the gradient preparation, beginning with the highest concentrated solution. DNA-lipid mixture after detergent removal was added to the top and centrifuged at 55 000 rpm for 6 h at 4 °C. After centrifugation, fractions were collected and analyzed with 6% SDS-PAGE analysis. The native-PAGE gel and running buffer described above were supplemented with 0.1% SDS, both of which were supplemented with 100 mM NaCl. The gel was pre-run at 100 V for 30 min at 15 °C before loading with the sample volume of 18 µL from each fraction with 2 µL of 10× loading buffer (50% Glycerol, 100 mM Tris HCl (pH 8.0), 0.1% SDS). The electrophoresis was performed at the same condition for 1 h. The gel was first imaged to identify the lipid fractions with Rhodamine signal and then stained with SYBR^TM^ Gold for DNA signal.

### Size exclusion chromatography

SEC was performed by JASCO HPLC system using either Superose^TM^ 6 Increase 10/300 GL column (Cytiva) at a flow rate of 0.5 mL/min or 3.2/300 GL column (Cytiva) at 0.04 mL/min and eluted with the buffer (10 mM HEPES (pH 7.4), 150 mM NaCl, 5 mM MgCl_2_). The eluted fractions were pooled with 30 kDa or 10 kDa MWCO Amicon filters according to the manufacturer’s protocol and kept at 4 °C until use.

### SDS-PAGE for protein analysis

SDS-PAGE for protein purified from DGUC and SEC analysis was prepared as follows: the running gel (15%) and stacking gel (5%) contain the required volume of 40% acrylamide/bis solution (29:1), 0.39 M Tris (pH 8), 0.1% SDS, 0.1% TEMED, and 1% APS. The gel was pre-run for 90 min prior to the sample loading using 1× SDS-PAGE running buffer (Nippon Genetics). The samples were mixed with the loading buffer without β-Mercaptoethanol, loaded along with the unstained and per-stained Precision Plus Protein^TM^ ladders (Bio-rad) and run for 2 h. The protein staining was performed with Sypro^TM^ Ruby Protein Gel stain (Thermo Fisher Scientific) according to the manufacturer’s protocol. The protein gel was visualized with iBright^TM^ CL 1500 Imaging System (Thermo Fisher Scientific).

### AFM imaging

AFM imaging was performed using a BIXAM microscope (Olympus, Tokyo, Japan). A drop (2 µL) of the sample in the buffer was deposited onto the freshly cleaved mica surface and incubated for 3 min. After the incubation, 2 µl of 0.05% 3-aminopropyltriethoxysilane was further deposited to the surface and the solution was incubated for 3 min. The surface was then rinsed with 10 µL of buffer composed of 10 mM Tris-HCl (pH 7.5), 0.1 mM EDTA, and 15 mM MgCl2. The micae surface was scanned in ∼120 µL of the buffer using a small cantilever (9 µm long, 2 µm wide, and 130 nm thick) with a spring constant of 0.1 N m−1 and a resonant frequency of 300–600 kHz in water (USC-F0.8-k0.1-T12; Nanoworld, Neuchâtel, Switzerland). AFM imaging was performed in an air-conditioned, temperature-controlled room at 25 °C.

### TEM imaging

TEM imaging was performed using a Tecnai G2 Spirit (FEI, United States). Carbon-coated TEM grids (Okenshoji, Japan) underwent plasma treatment for 10 s. A 15 nM sample, purified by SEC and concentrated by Amicon, was mixed with 1% trehalose in a buffer on the grid, to obtain 10 nM sample and 0.3% trehalose as a final concentration. The grid was incubated for 10 min at 25 °C, then followed by removing excess solution with filter paper. Next, the sample was stained with aqueous uranyl acetate solution (1 %(w/v)) dissolved in a buffer for 5 min at 25 °C, and the water was removed with filter paper. The grid was visualized at 120 kV acceleration voltage.

### Atomistic Molecular Dynamics Simulations

Two circular DNA nanodisc (147 bp) was generated using the CGeNArateWeb server^61^. Subsequently, PyMOL^62^ was used to refine the double-decker structure, and fifty-eight nucleotides were functionalized with heptyl alkyl chains. A pre-equilibrated lipid bilayer composed of DMPC and POPC (85%) and DMTAP and DOTAP (15%) was obtained from CHARMM-GUI^63–65^ and combined with the DNA double-decker. Lipids located outside the DNA boundary of the DNA were removed using the MDAnalysis library^66,67^, resulting in a final nanodisc containing either 477 lipids (400 DMPC and 77 DMTAP) or 448 lipids (371 POPC and 77 DOTAP), and a clash-free structure suitable for subsequent energy minimization and equilibration.

To optimize computational efficiency and minimize the required solvent volume, a compact simulation box (22 × 22 × 14 nm³) was employed, and a minimum distance of 1.2 nm was maintained between the DNA nanodisc and the box boundaries. To prevent DNA nanodisc from diffusing toward the periodic boundaries, flat-bottom position restraints with a layer geometry were applied to eight backbone phosphorus atoms. These restraints defined a 6.0 nm unrestrained slab (±3.0 nm) along the normal nanodisc (z-axis). Within this region, the DNA scaffold was permitted free and unbiased motion, while a harmonic force constant (1000 kJ mol⁻¹ nm⁻²) was applied only when the atoms moved beyond the ±3.0 nm threshold from their reference coordinates. Then the system was neutralized, and NaCl and MgCl₂ were added to final concentrations of 200 mM and 5 mM, respectively.

To maintain structural stability, base-pairing interactions were governed by piecewise linear/harmonic distance restraints (GROMACS function type 10) with a force constant of 1000 kJ mol⁻¹ nm⁻². These restraints were applied to specific base-pairing atoms (N1 of purines and N3 of pyrimidines), as unrestrained simulations exhibited significant disruption of hydrogen bonding under the CHARMM force field, consistent with previous studies reporting reduced base-pair stability in unrestrained DNA simulations^68^.

Before equilibration, the system underwent a two-stage energy minimization using the steepest descent algorithm. In the first stage, no bond constraints (e.g., LINCS) were applied, and the systems converged (Fmax < 1000 kJ mol⁻¹ nm⁻¹). In the second stage, constraints were applied to bonds involving hydrogen atoms using the LINCS algorithm, consistent with the CHARMM36 force field, and the system converged (Fmax < 1000 kJ mol⁻¹ nm⁻¹). The minimized structure obtained from the second stage was used as the reference structure (-r) for position restraints during subsequent equilibration.

After minimization, the resulting clash-free and stabilized structure was used for a six-step equilibration protocol adapted from CHARMM-GUI nanodisc setups^63–65,69^, consisting of three initial 0.125 ns steps with a 1 fs time step followed by three 0.5 ns steps with a 2 fs time step (totaling 1.875 ns). Position restraints on the DNA and protein backbone, side chains, and lipids were gradually reduced (DNA backbone: 4000 → 50 kJ mol⁻¹ nm⁻²; side chains: 2000 → 0 kJ mol⁻¹ nm⁻²; lipids: 1000 → 0 kJ mol⁻¹ nm⁻²), along with dihedral restraints (1000 → 0 kJ mol⁻¹ rad⁻²).

Molecular dynamics simulations were performed using GROMACS (2024.4)^70^ with the CHARMM36 force field and CUFIX corrections^71^ and the TIP3P water model. Heptyl chain parameters were derived from CGenFF^72–74^, with missing bonded terms assigned by analogy using Gromologist^75^. The system was integrated using the leap-frog algorithm with a 2 fs time step for a total production time of 200 ns. Simulations were conducted at 310 K (above lipid phase transition temperature to ensure fluid phase) for DMPC and DMTAP and 298K for POPC and DOTAP.

The temperature was controlled using the velocity-rescaling (v-rescale) thermostat^76^. Pressure was maintained at 1.0 bar using the C-rescale barostat^77^ with isotropic coupling. Long-range electrostatic interactions were treated using the Particle Mesh Ewald (PME) method^78^ with a real-space cutoff of 1.2 nm. Van der Waals interactions were calculated using a force-switch modifier between 1.0 and 1.2 nm. All bonds involving hydrogen atoms were constrained using the LINCS algorithm^79^. Trajectories were saved every 100 ps. The resulting trajectories were corrected using the MDVWhole algorithm^80^ to remove periodic boundary artifacts.

### Analysis

The stereochemical integrity of the modified phosphorothioate centers was verified using an in-house Python pipeline incorporating the RDKit library^81^, which assigned the R and S configurations of each modified residue based on their molecular structures, ensuring consistency with the intended chirality.

---

**Supplementary Table 1.**
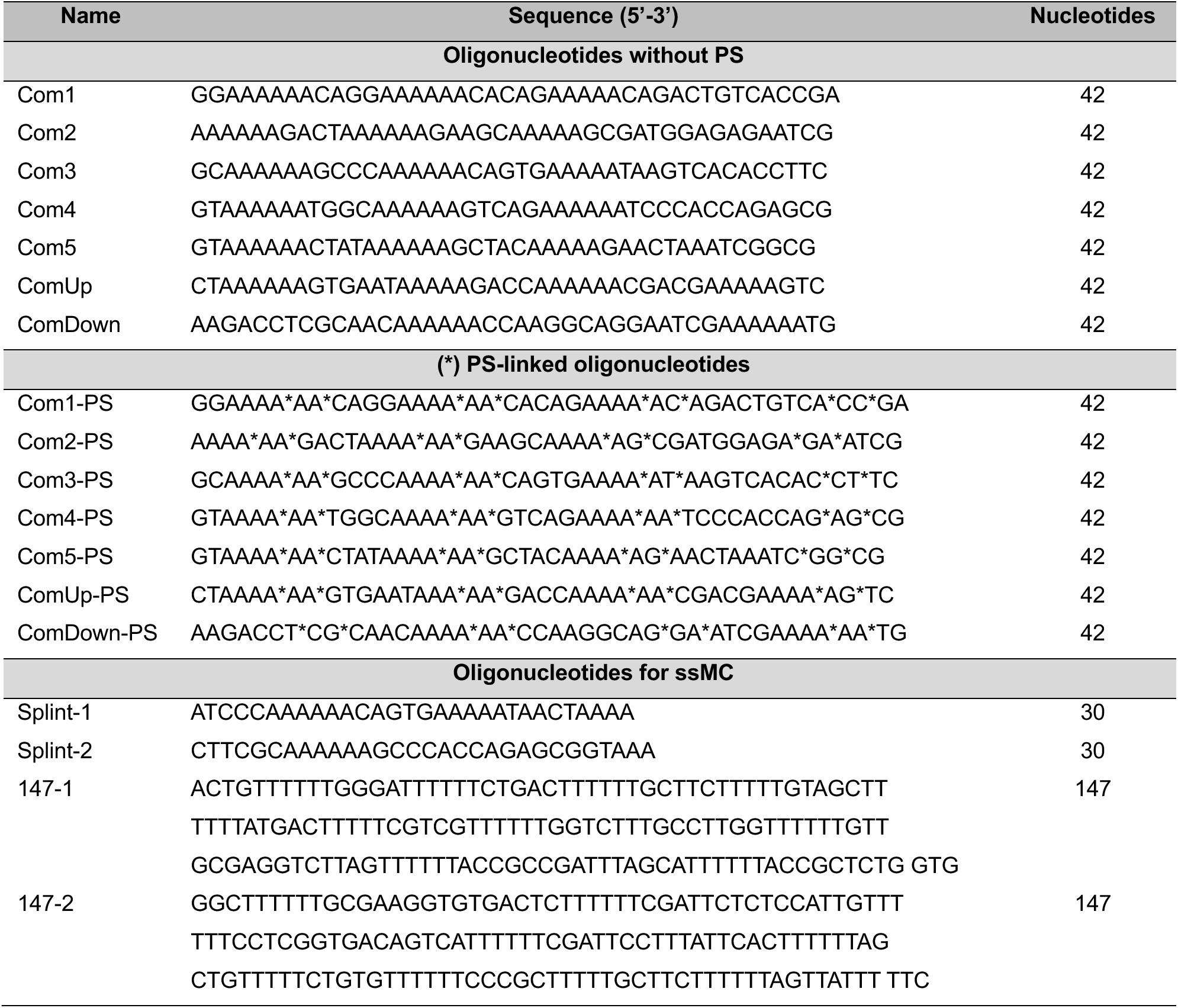
List of oligonucleotides used in this study.

**Figure S1.**
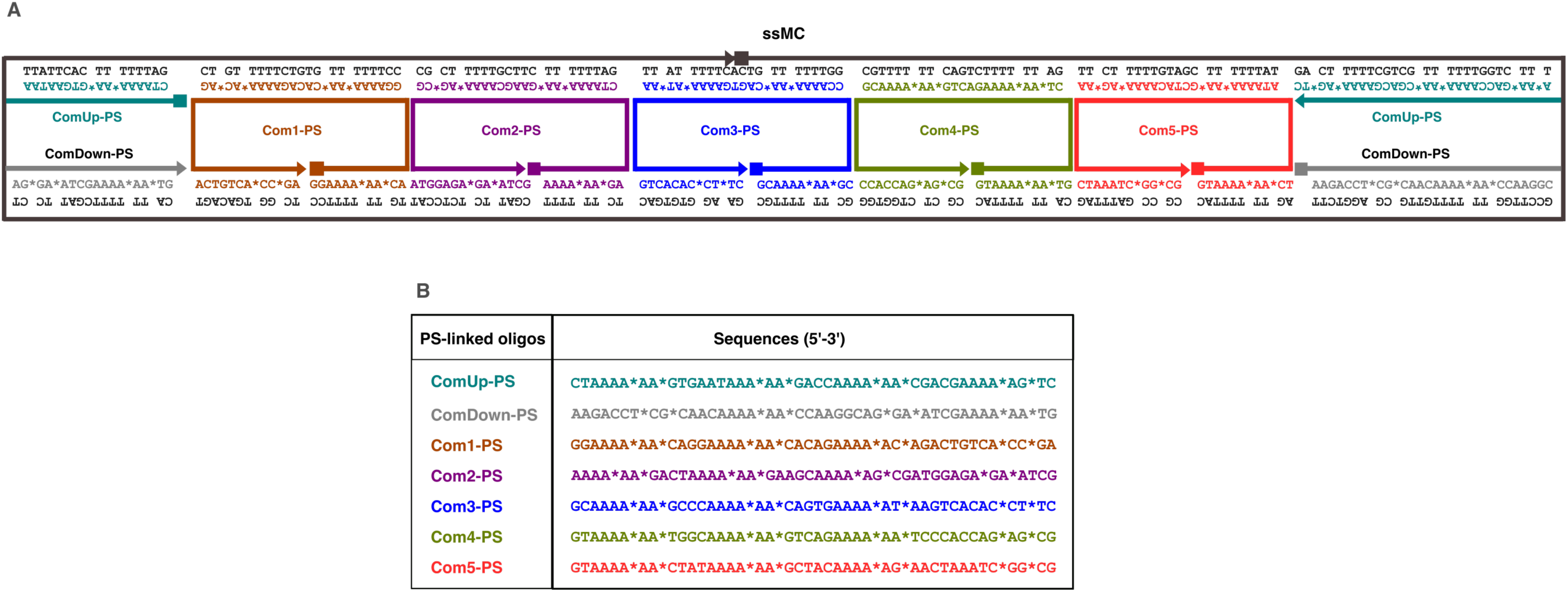
(A) Design map of the DDring denoting the crossovers between the staples and the ssMC. (B) Sequences of ssDNA oligonucleotides used as staples for assembling DDring, (also shown in Table S1). (*) denotes the location of PS.

**Figure S2.**
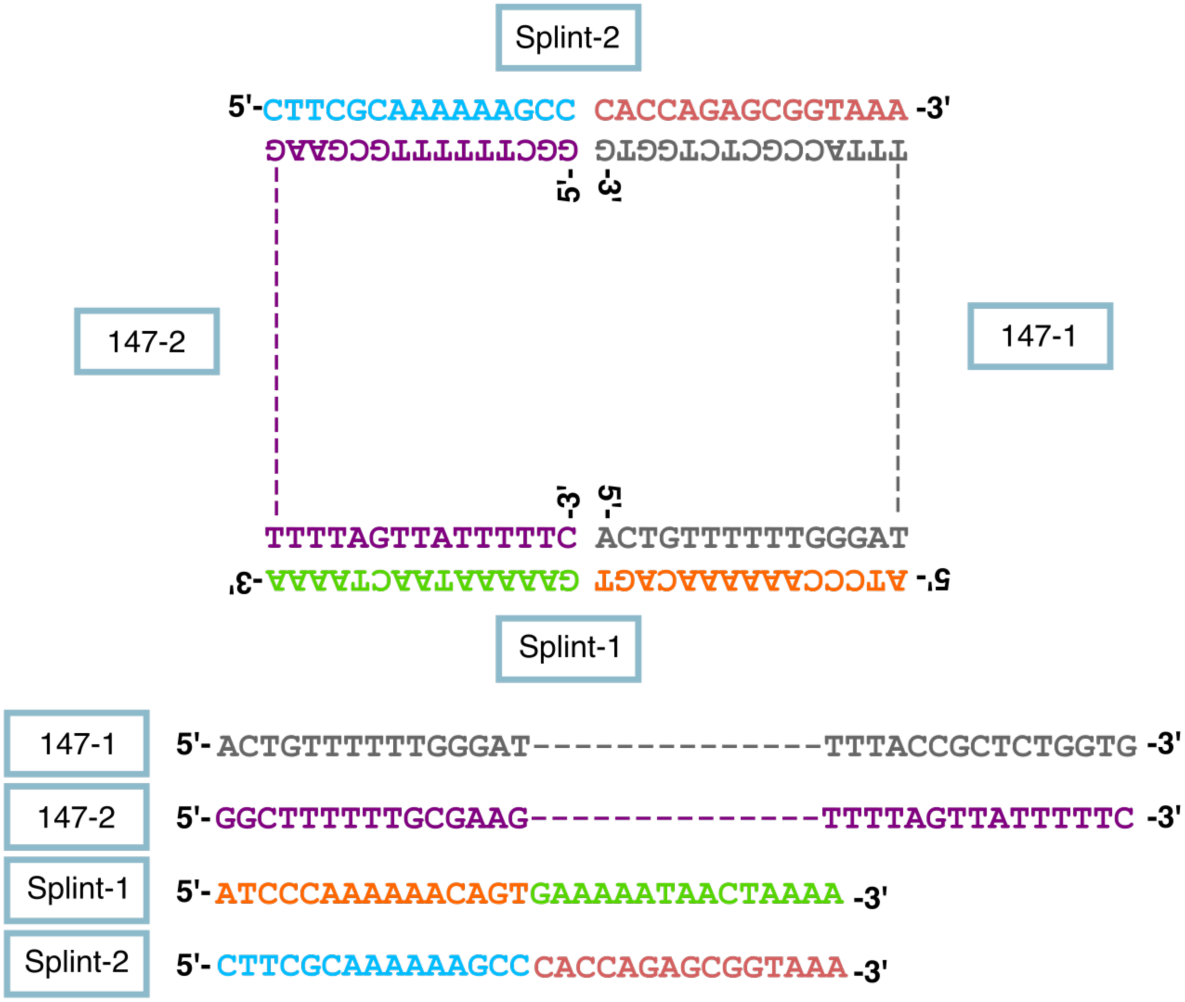
Sequence map of oligonucleotides in splint ligation to generate ssMC. Full sequences of two 147-mer sequences are listed in table S1.

**Figure S3.**
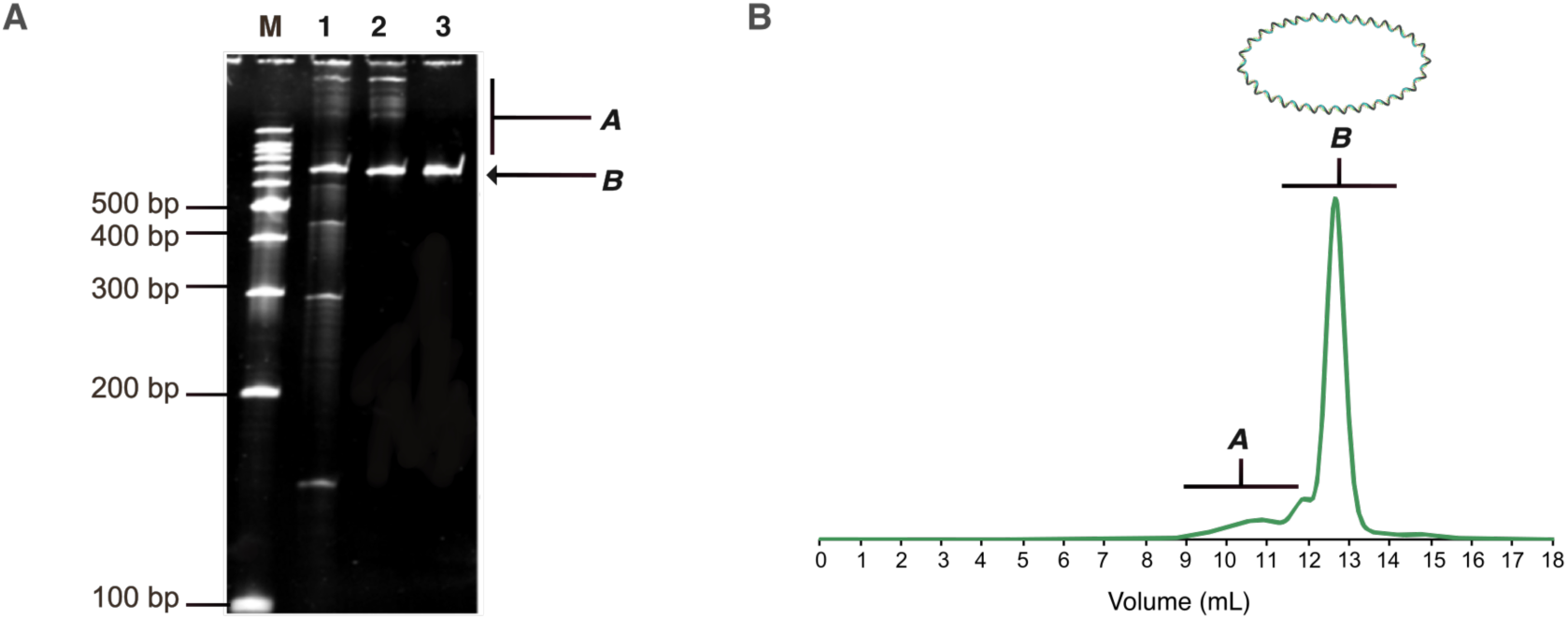
Denaturing PAGE analysis of splint-ligated ssMC with 5% gel for samples (1) before and (2) after exonuclease treatment, and (3) after SEC purification. (B) SEC profile shown in figure 1C is repeated here to denote the pattern of different fractions, A and B. The ligation product before exonuclease contains the remaining splints and non-ligated linear DNAs. Treatment with exonuclease I and III eliminates the linear DNAs while dimeric ssMC can only be eliminated by SEC. M = 100 bp ladder marker.

**Figure S4.**
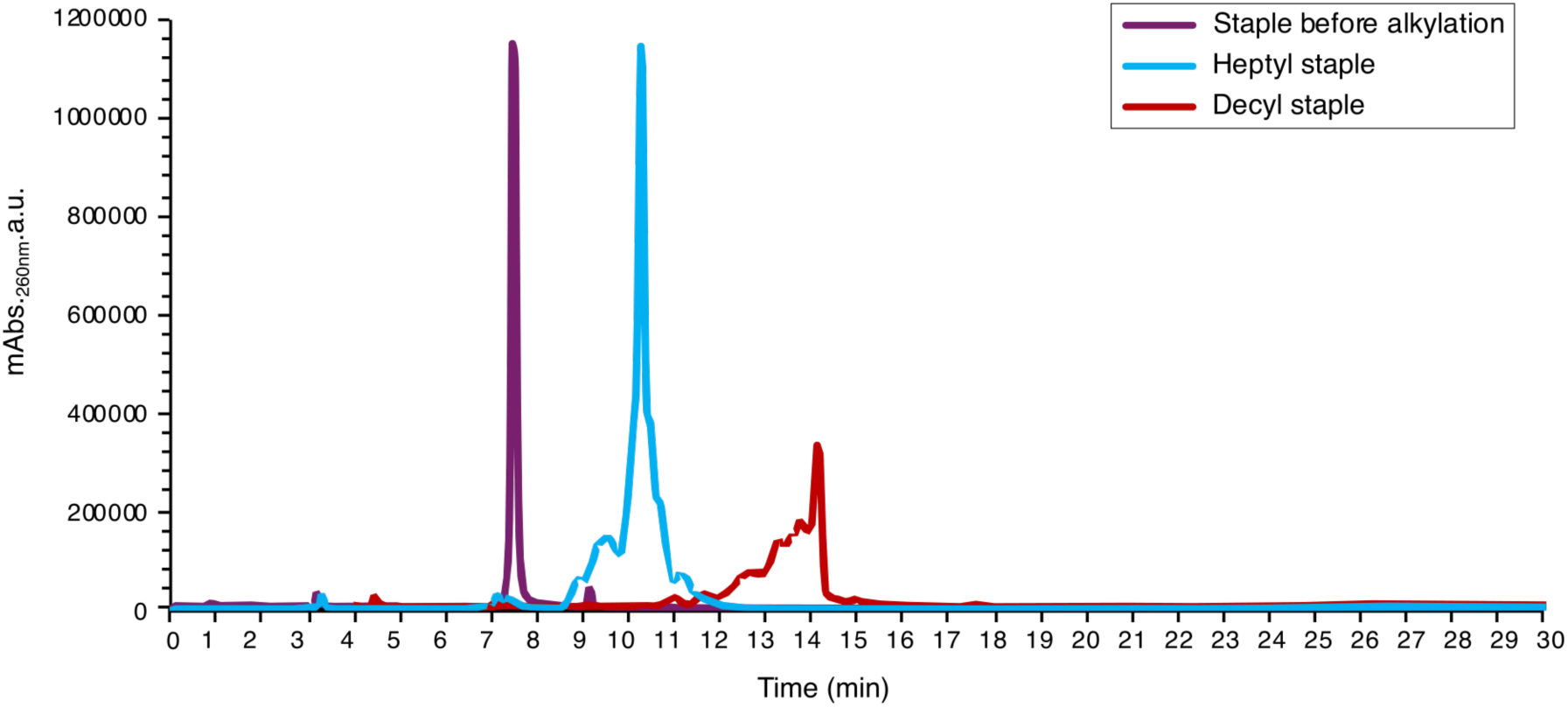
RP-HPLC analysis of PS-linked oligos. Comparison of peak intensity between staple before alkylation and after alkylation using heptyl and decyl moieties. Elution time takes longer time with the more hydrophobic samples.

**Figure S5.**
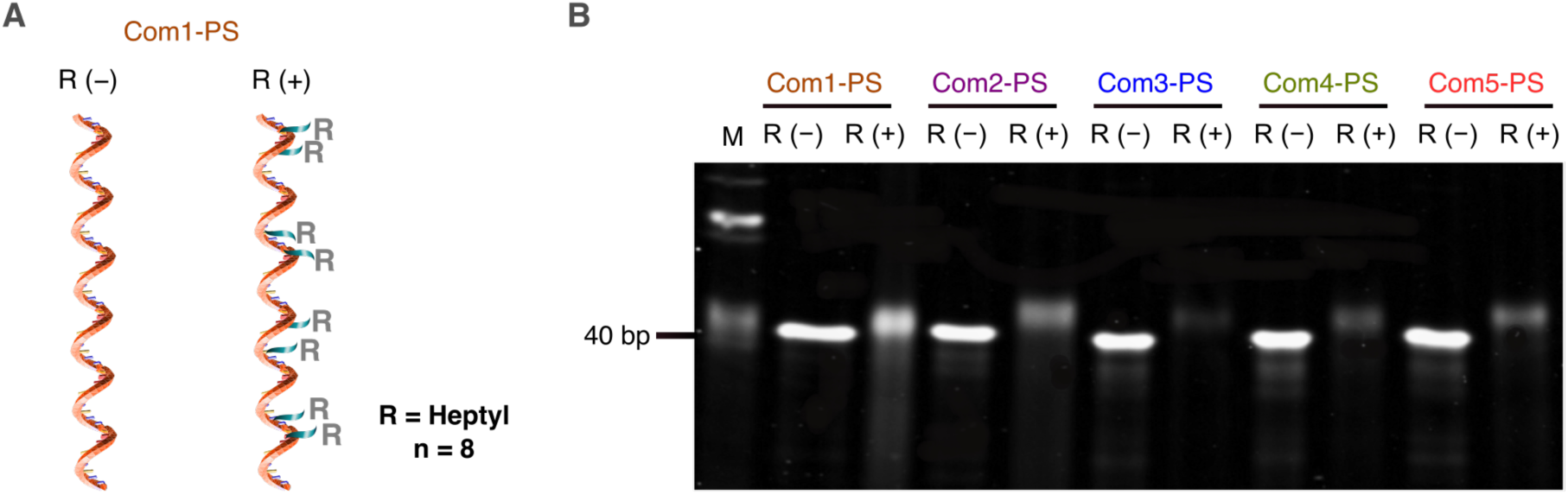
(A) PS-linked oligonucleotide with and without alkyl moieties. Each 42-mer oligo possesses 8-PS modifications. (B) Denaturing PAGE gel (16%) of 5 oligos out of 7, Com1-5-PSs, before and after alkylation. Marker = 20 bp ladder.

**Figure S6.**
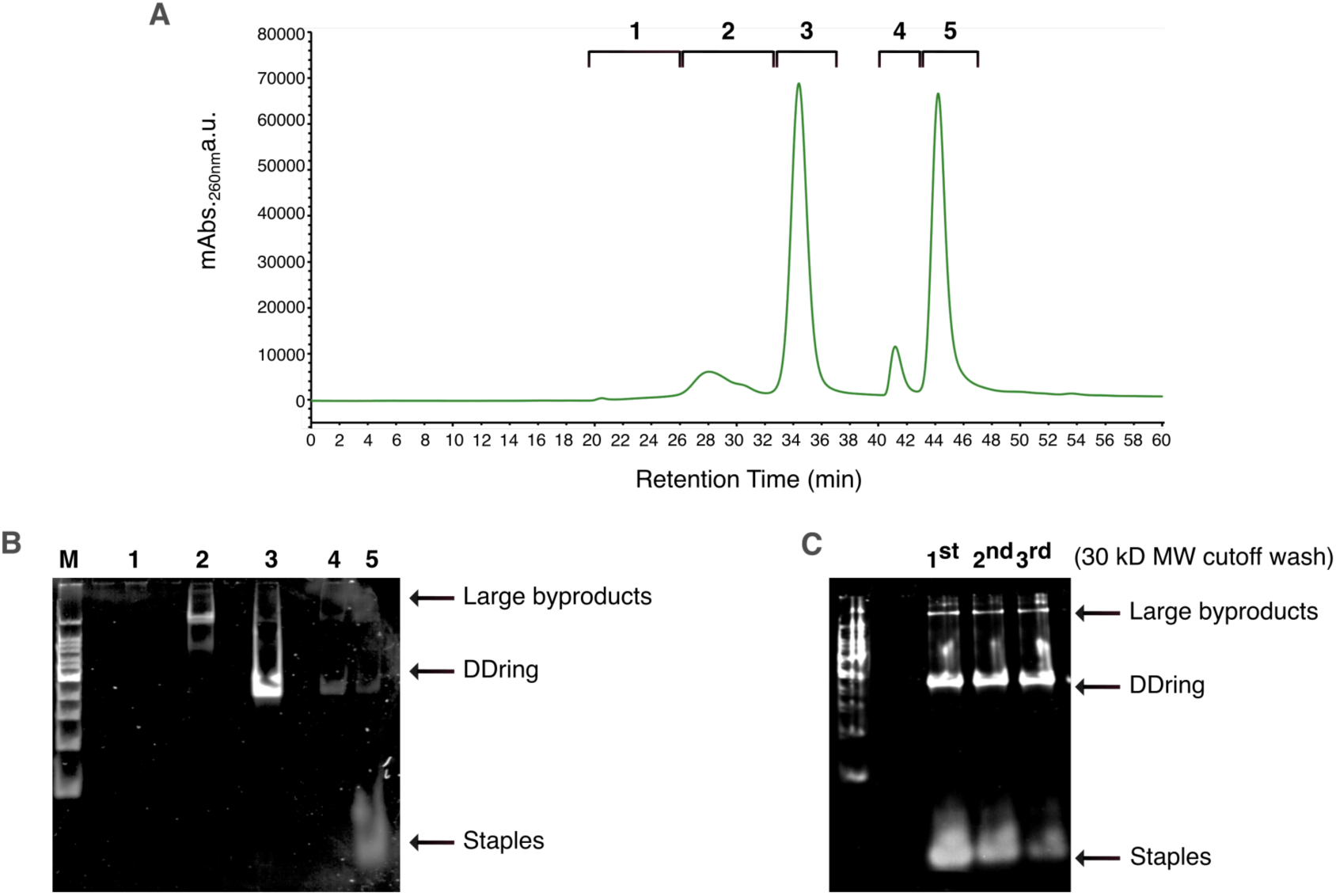
(A) SEC chromatogram of DDring before washing the annealing sample off using 30 kD MW cutoff Amicon filtration column that remove excessed staples. (B) Native-PAGE analysis (4%) of the collected fractions: 1, 2, 3, 4 and 5 from SEC chromatogram in A. (C) Native-PAGE analysis (4%) of DDring after washing multiple times using 30 kD (1st, 2nd and 3rd).

**Figure S7.**
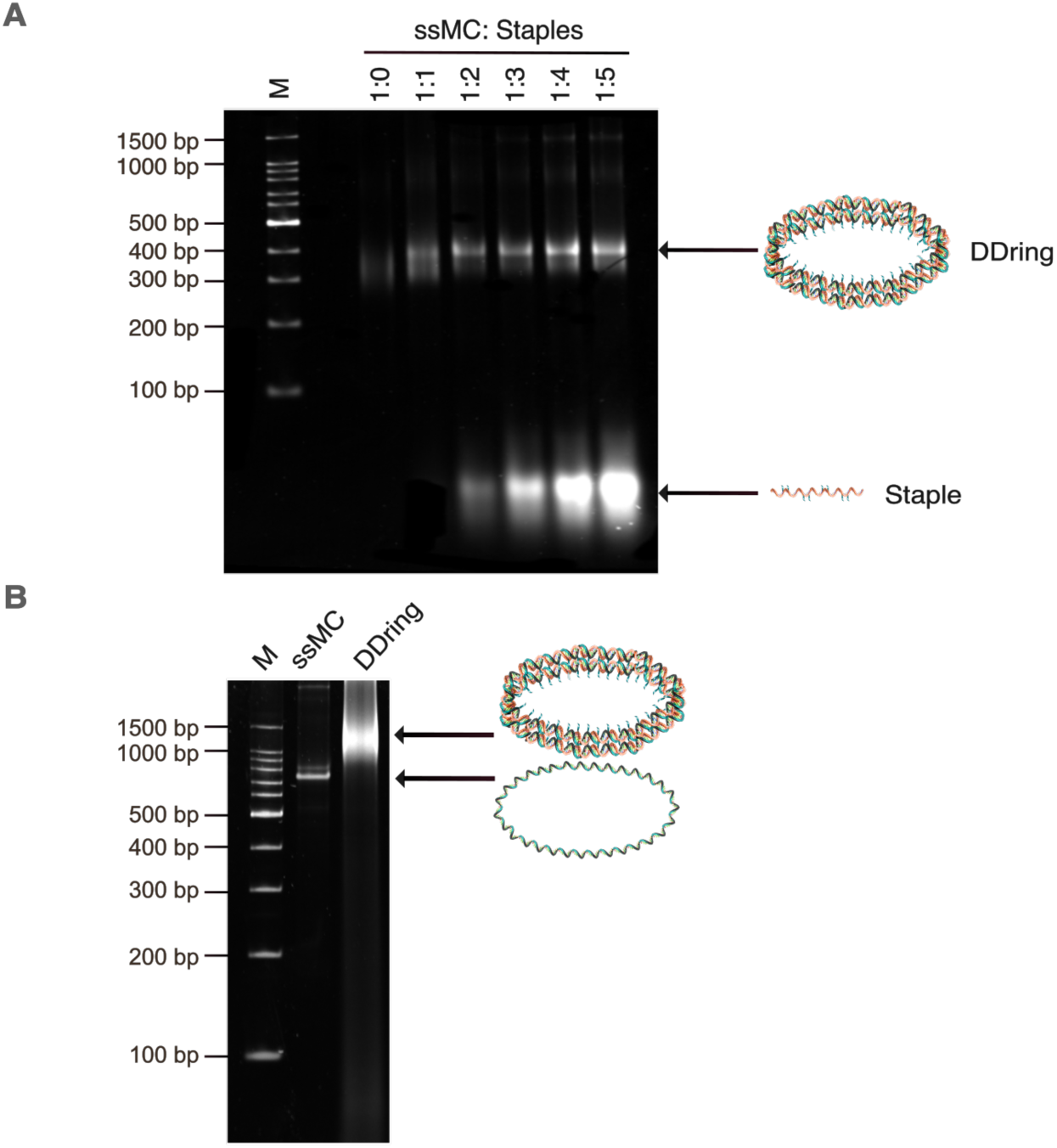
(A) Native PAGE (4%) analysis of the yield of DD ring with different ssMC to staples ratios, from left to right 1:0.5, 1:1, 1:2, 1:3, 1:4, 1:5. (B) Native-PAGE (5%) analysis of DDring with heptyl moieties (n = 56) in comparison to ssMC. Marker = 100 bp ladder DNA.

**Figure S8.**
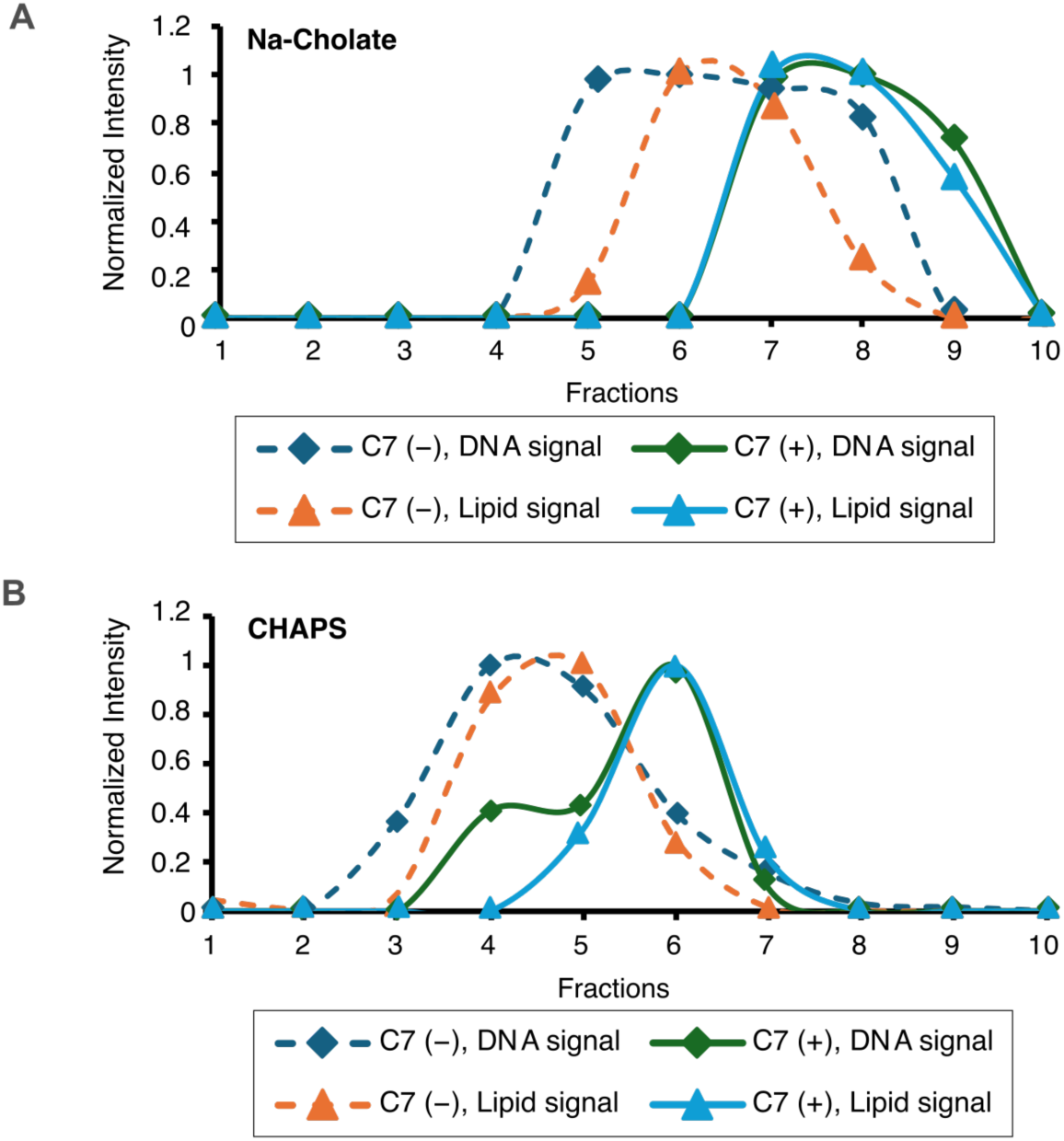
(A) Comparison of the lipid and DNA intensity after the density gradient ultracentrifugation (DGUC). DMPC:DMTAP:Rhod-PE (89:10:1) lipids are solubilized in the sodium cholate detergent, mixed with DDrings (Alkyl −/ +) and subjected to DGUC after detergent removal. (B) SDS-PAGE analysis between alkyl and non-alkyl DDrings after DGU using DMPC:DMTAP:Rhod-PE (89:10:1) lipids and CHAPS detergent. Rhodamine-PE containing lipids (red) signal in the gel was first imaged prior to the staining with Sybr-Gold to visualize DNA signal. Fractions (1-10) were taken by pipetting from top to bottom representing from less to more Iodixanol intensity. (C) Comparison of the lipid and DNA intensity from B.

**Figure S9.**
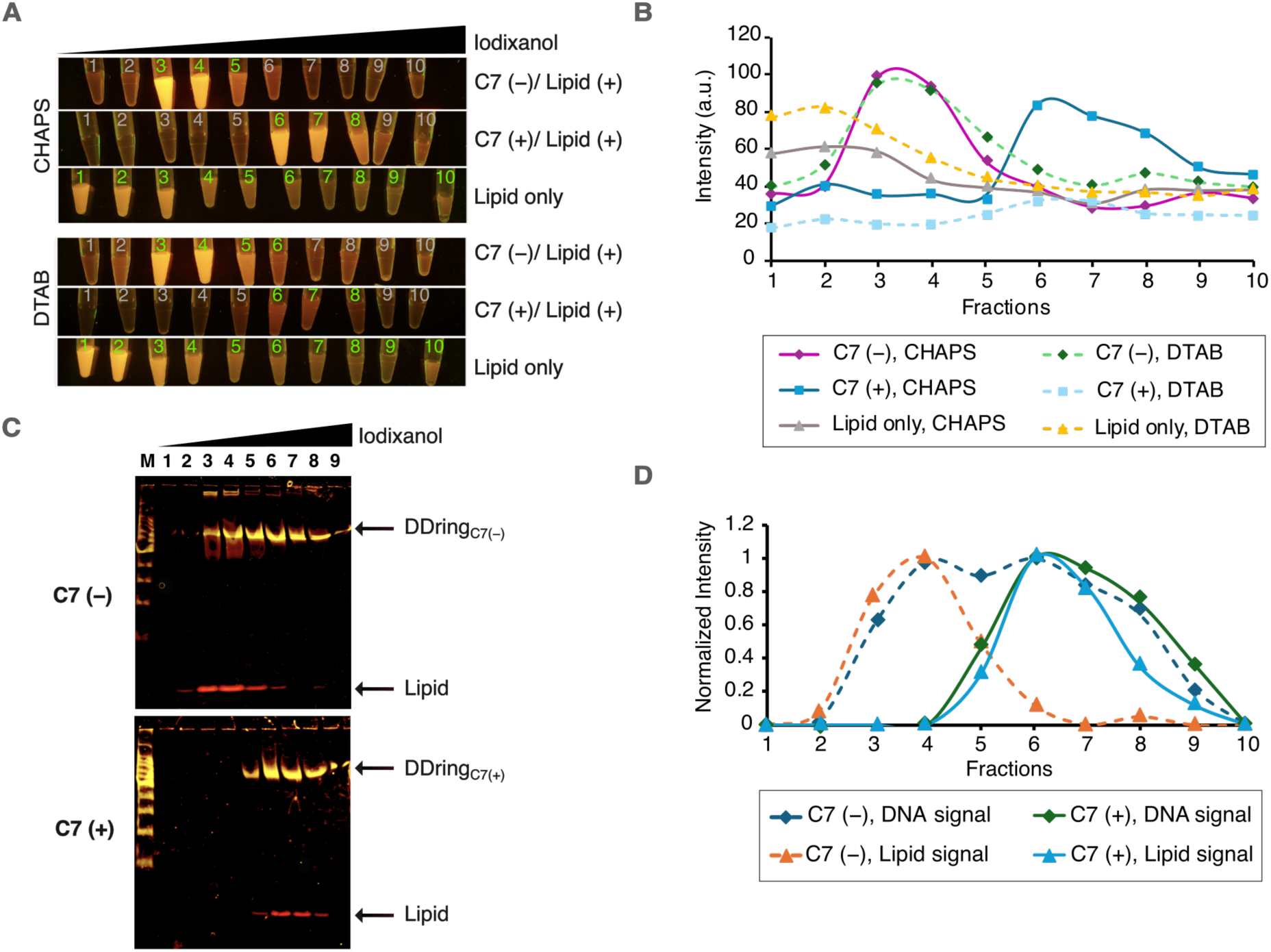
SDS-PAGE analysis between alkyl and non-alkyl DDrings after DGU using POPC:DOTAP:Rhod-PE (89:10:1) lipids and DTAB detergent. Rhodamine-PE containing lipids (red) signal in the gel was first imaged prior to the staining with SYBR Gold to visualize DNA signal. Fractions were taken from top to bottom by pipetting. Comparison of the lipid and DNA intensity were shown in figure 2 along with CHAPS detergent batch.

**Figure S10.**
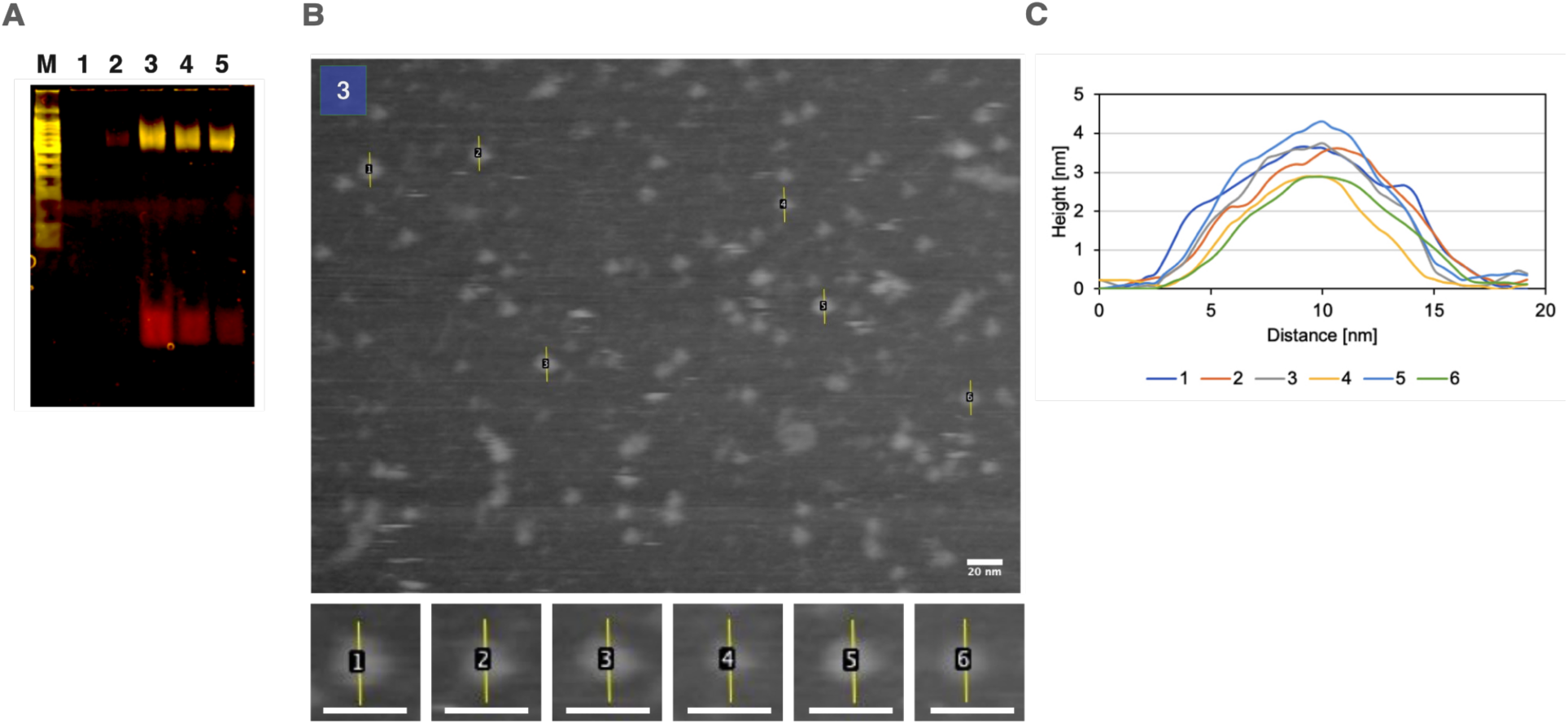
(A) SDS-PAGE gel (6%) of alkyl DDrings after DGU using POPC:DOTAP:Rhod-PE (89:10:1) lipids and CHAPS detergent. (B) Fraction-3 was subjected to AFM imaging. (C) Height and distance plot of the selected areas (1-6) from B.

**Supplementary Movie 1.**
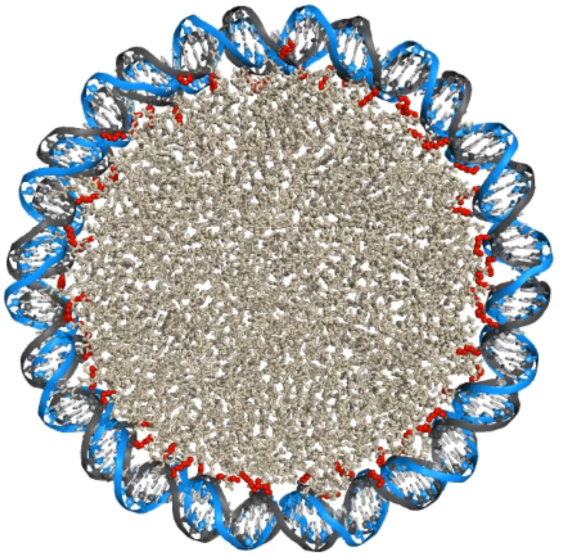
All-atom molecular dynamics simulation of the DNA nanodisc (DMPC/DMTAP). Top view over a 200 ns trajectory. The lipid environment is shown as light gray, with heptyl groups represented as red spheres and heptyl-modified DNA strands in dark gray. The DNA scaffold is shown in light blue. The snapshot shown is taken from the first frame of the 200 ns trajectory.

**Supplementary Movie 2.**
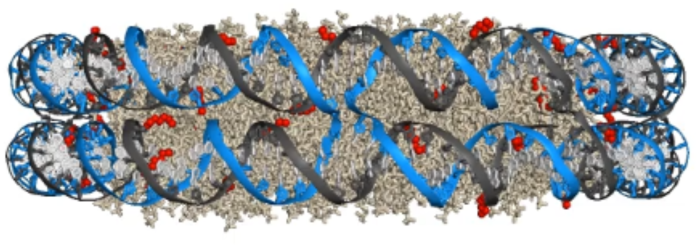
All-atom molecular dynamics simulation of the DNA nanodisc (DMPC/DMTAP). Side view over a 200 ns trajectory. The lipid environment is shown as light gray, with heptyl groups represented as red spheres and heptyl-modified DNA strands in dark gray. The DNA scaffold is shown in light blue. The snapshot shown is taken from the first frame of the 200 ns trajectory.

**Supplementary Movie 3.**
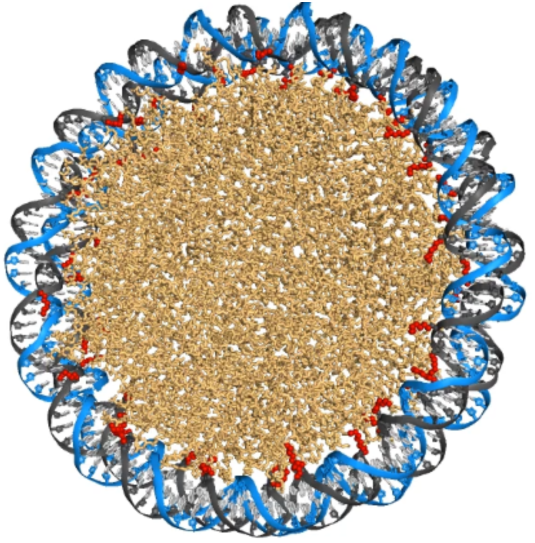
All-atom molecular dynamics simulation of the DNA nanodisc (POPC/DOTAP). Top view over a 200 ns trajectory. The lipid environment is shown as light orange, with heptyl groups represented as red spheres and heptyl-modified DNA strands in dark gray. The DNA scaffold is shown in light blue. The snapshot shown is taken from the first frame of the 200 ns trajectory.

**Supplementary Movie 4.**
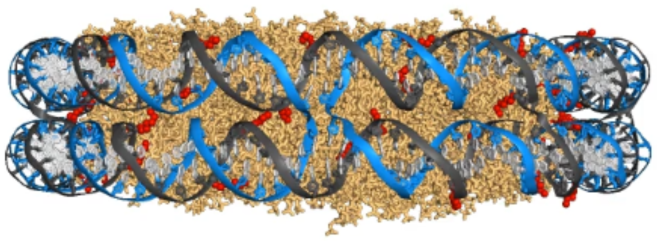
All-atom molecular dynamics simulation of the DNA nanodisc (POPC/DOTAP). Side view over a 200 ns trajectory. The lipid environment is shown as light orange, with heptyl groups represented as red spheres and heptyl-modified DNA strands in dark gray. The DNA scaffold is shown in light blue. The snapshot shown is taken from the first frame of the 200 ns trajectory.

**Supplementary Movie 5.**
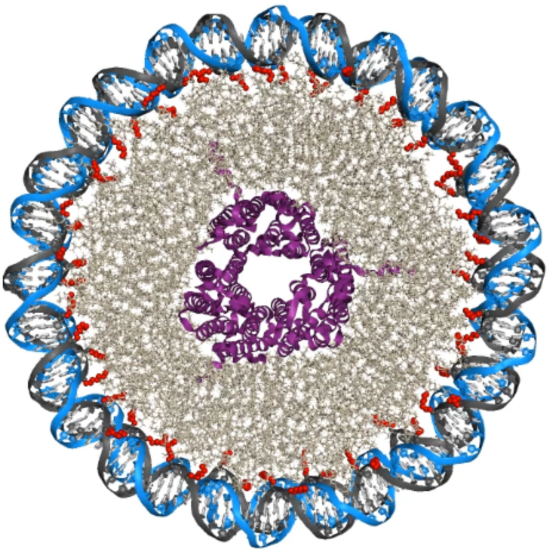
All-atom molecular dynamics simulation of the DNA nanodisc (DMPC/DMTAP) with membrane protein. Top view over a 200 ns trajectory. The lipid environment is shown as light gray, with heptyl groups represented as red spheres, heptyl-modified DNA strands in dark gray and the membrane protein (bR) in purple. The DNA scaffold is shown in light blue. The snapshot shown is taken from the first frame of the 200 ns trajectory.

**Supplementary Movie 6.**
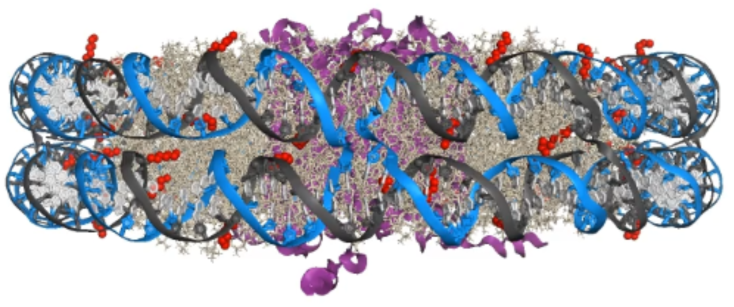
All-atom molecular dynamics simulation of the DNA nanodisc (DMPC/DMTAP) with membrane protein. Side view over a 200 ns trajectory. The lipid environment is shown as light gray, with heptyl groups represented as red spheres, heptyl-modified DNA strands in dark gray and the membrane protein (bR) in purple. The DNA scaffold is shown in light blue. The snapshot shown is taken from the first frame of the 200 ns trajectory

## Notes

### Competing Interest Statement

The authors have declared no competing interest.

